# The Arabidopsis UMAMIT30 transporter contributes to amino acid root exudation

**DOI:** 10.1101/2025.08.23.671809

**Authors:** Israel D. K. Agorsor, Pramod Khadka, Cristian H. Danna

## Abstract

**Background:** Root exudation is an important trait that enables plants to shape their interactions with soil-borne organisms. Amino acids present in root exudates play important roles in bacterial chemotaxis, bacterial metabolism, and root colonization, contributing to plant nutrition and health. Notwithstanding the importance of amino acids in shaping the rhizosphere microbiome, the identities of the plant amino acid transporters that mediate their root exudation have remained elusive.

**Results:** Here, we report that the Arabidopsis UMAMIT30 transporter, robustly expressed in root and shoot tissues, significantly contributes to amino acid root exudation. *umamit30* loss-of-function mutants were compromised for amino acid root exudation as shown by the low concentration of amino acids, particularly glutamine, recovered from root exudates compared to wild-type plants. Amino acid quantification, as well as uptake and secretion assessments using radiolabelled glutamine, revealed that the shoots of *umamit30* accumulate amino acids and have a reduced capacity to secrete glutamine, impacting root exudation.

**Conclusions:** Our results identify UMAMIT30 as a broadly specific amino acid exporter strongly expressed in Arabidopsis vasculature. Loss-of-function mutants displayed reduced amino acid levels in root exudates, with significant drops in glutamine and asparagine among others, yet exhibited no detectable growth defects. UMAMIT30 disruption led to elevated shoot amino acid content and reduced glutamine efflux from shoots, suggesting a role in phloem uploading rather than root exudation alone. Despite decreased levels of root exuded amino acids, beneficial *Pseudomonas* interactions and plant-growth-promotion remained unaffected.

## Introduction

Plant roots mediate plant―environment interactions by anchoring plants to their growth substrate, acquiring nutrients and water, and exuding plant metabolites that shape the microbial composition of the narrow region of the soil around the roots, i.e., the rhizosphere [1]. Plants can shape their rhizosphere in ways that impact plant nutrition, plant growth, and plant health, leading to the notion that the rhizosphere is an extended phenotype of the root [2]. Plant root exudates usually consist of primary metabolites such as sugars, amino acids (AAs), and organic acids, as well as secondary metabolites [3, 4, 5, 6]. These plant-made molecules not only serve as nutrients but also as signalling molecules that attract or repel microbes and promote microbial competition [7].

In the plant growth-promoting rhizobacterium *Bacillus subtilis*, AAs serve as ligands for many of the chemoreceptors encoded in the genome. Thus, root exudate components such as AAs may facilitate root colonization by *B. subtilis* [8] and other microbes. Notably, some rhizosphere microbes harbouring mutations in genes involved in AAs assimilation and catabolism are unable to colonize roots efficiently [9, 10]. Additionally, the expression of genes that regulate competence and sporulation in *B. subtilis* is increased by rhizosphere AAs [11], suggesting that rhizosphere AAs in part govern the assemblage of the microbiome. When AAs are available in the rhizosphere, microbes outcompete plant roots at using them [12]. In addition, rhizobacteria belonging to the *Pseudomonas* and *Micromonospora* genuses prefer AAs to sugars as carbon and nitrogen sources [13, 14, 15]. Also, in nutrient-limiting environments, bacterial survival may be enabled by mutations that allow for increased AA catabolism [16]. Furthermore, bioinformatics analysis suggested that, between the rhizosphere-dwelling bacterium *P. putida* KT2440 and its close relative *P. aeruginosa*, an opportunistic human pathogen, *P. putida* KT2440 encodes twofold more genes for AA transporters than does *P. aeruginosa* [17]. These findings imply that rhizobacteria adapt to AA concentration changes in the rhizosphere to increase their fitness.

Plants obtain nitrogen from the soil as inorganic salts, which are converted into AAs in the roots or mature leaves. These AAs are then transported to distal tissues and organs to support plant growth. For example, AAs are typically transported from leaves after synthesis to heterotrophic sink tissues such as developing leaves, apical meristems, and stem-localized cortical cells, as well as seeds, fruits, and roots [18]. In Arabidopsis, root exudation is temporally regulated and dependent on the developmental stage, with the exudation of AAs increasing over time [19]. Furthermore, the expression of AA metabolism genes in soil microbes, for example, follows the pattern of AA accumulation in root exudates [19, 20], highlighting what is potentially an outsized role for root-exuded AAs in shaping root–microbe interactions.

Notwithstanding their prevalence in root exudates, the mechanisms by which AAs are secreted from roots or taken up back into the root from the rhizosphere remained unknown until recently. Transporter of AAs with uptake activity in Arabidopsis were first described in the 1990s [21, 22, 23, 24]. Among others, the LHT1 (Lysine-Histidine Transporter-1) transporter plays a key role taking up AAs against a concentration gradient. Furthermore, LHT1 plays an important role modulating the concentration of glutamine and other AAs in root exudates. The accumulation of an excess of glutamine in *lht1* root exudates enhances chemotaxis and root colonization but compromises the ability of the plant-growth promoting bacteria *Pseudomonas simiae* strain WCS417r (hereinafter *Ps* WCS417r) to promote plant growth [25]. These data suggest that the amino acidic composition of the root exudates modulates bacterial metabolism, impacting an otherwise beneficial interaction between Arabidopsis roots and plant-growth promoting bacteria [25]. How AAs are secreted from plant cells was poorly understood until more recently. Although early physiological studies revealed AA export activity in plant cells, the exact molecular identities of the transporters involved have long remained unknown [26]. Data presented in several recent studies, however, showed that a group of transporters sharing high identity with the *Medicago truncatula Nodulin-21* gene, later named UMAMIT transporters (<u>U</u>sually <u>M</u>ultiple <u>A</u>mino acids <u>M</u>ove In and out <u>T</u>ransporters), are indeed facilitator transporters that exhibit AA export capabilities [27, 28, 29, 30, 31, 32, 33]. In particular, UMAMIT14 has been shown to contribute to unloading AAs from the root phloem into the root stele apoplast, a crucial step toward root-to-soil AA secretion [29]. The compromised secretion of root AA in *umamit14* roots is further aggravated in the double mutant *umamit14-umamit18* [29], suggesting that several UMAMITs may participate in the secretion of AAs at different steps in different tissues. Further supporting the role of UMAMIT facilitators in root exudation, UMAMIT04, UMAMIT06, UMAMIT37, and UMAMIT42 are expressed in epidermal root cells and root hairs [34], strategically positioning them to contribute to executing the final steps in root-to-soil AA secretion.

In this work, we present evidence showing that UMAMIT30 plays an important role in transport events leading to the upload of AAs to the phloem in leaves and the exudation by the roots. Albeit lower than wild-type levels, the concentration of AAs in *umamit30* root exudates did not impact either *in vitro* growth of the beneficial rhizobacterium *Ps* WCS417r, or its ability to promote plant growth.

## Materials and methods

### Plant materials and genetic analysis of *umamit30* mutants

Wild-type *Arabidopsis thaliana* Col-0 plants (ABRC accession number CS1092) were grown from seeds, and the *umamit30* mutants were derived from the Col-0 background. For genetic analysis of mutants, genotyping was performed as follows: insertion alleles for the *UMAMIT30* gene were identified in the Salk Arabidopsis T-DNA Insertion Mutant Collection [35]. The *umamit30-1* mutant corresponds to Salk_140547C, and genotyping was performed via LBb1.3 and the following gene-specific primers: forward, 5’- GCATTGAAGCGTACCAAAGAC-3’, and reverse, 5’-TTCTTGATGGAGGCATCAATC-3’ (LBb1.3+Reverse yields the *umamit30-1* insertion-specific product). The *umamit30-2* mutant corresponds to Salk_146977C, and genotyping was performed via LBb1.3 and the following gene-specific primers: forward, 5’-GAAATATTGCATCAAGCTCGC-3’, and reverse, 5’-CCTTCCACCGAGAAAAACAG-3’ (LBb1.3+Reverse produces the *umamit30- 2* insertion-specific product). The primers were used at a final concentration of 10 µM. A total volume of 20 µL was used for each reaction with Quick-Load® Taq 2x Master Mix (New England BioLabs) (10 µL), primer pairs (1 µL), genomic DNA (2 µL), and DEPC- treated water (7 µL). For genotyping each line, an additional control PCR was performed using genomic DNA, as well as a blank reaction in which the DNA template was replaced with DEPC-treated water. The PCR conditions are as indicated below:

Step 1: Initial denaturation at 94°C for 5 mins; Step 2: Denaturation at 94°C for 30 s; Step 3: Annealing at 55–56°C for 30 s; Step 4: Extension at 68°C for 1 min; Step 5: Final extension at 68°C for 5 mins. Steps 2-4 were repeated for a total of 35 cycles. Step 6: Hold at 12°C. The PCR products were resolved on a 1% (w/v) ethidium bromide-stained agarose gel run at 100 V. Gels were visualized via the Spectroline UV transilluminator SelectTM Series, and images were obtained via the AlphaImager 2200.

RT‒PCR: Roots/shoots/leaves harvested from wild-type and *umamit30* mutants as well as *umamit14* and *umamit05* mutants were frozen in dry ice. RNA was isolated via the use of TRIzol® Reagent (Fisher Scientific; Cat# 15596018.) and quantified with a NanoDrop- ND1000 spectrophotometer (Thermo Fisher Scientific, Inc.). Two micrograms of total RNA was used for 1st strand cDNA synthesis after DNaseI treatment. The 2 µg RNA was incubated together with 2 µL of random decamers and an appropriate volume of DEPC-treated water to a final volume of 15 µL at 70°C for 5 minutes and then immediately transferred to ice. The cDNA synthesis mixture containing 10 mM dNTPs (2 µL), Promega RNase inhibitor (1 µL), Promega M-MLV-RT (1 µL), 5x Promega M-MLV Buffer (5 µL) and DEPC-treated water (1 µL) was subsequently added to a final volume of 25 µL, which was subsequently incubated at room temperature for 1 minute and then at 37°C for 60 minutes. For PCR analysis, 2 µL of cDNA was used in a total reaction volume of 20 µL containing 10 µL of Quick-Load® Taq 2x Master mix, 1 µL of 5 µM primers (forward and reverse), and 7 µL of DEPC-treated water. The sequences of primers used for the RT‒ PCR analyses are shown in Supplementary Table S1. A “no template control” (i.e., blank) PCR was performed in which the cDNA template was replaced with DEPC-treated water. The PCR conditions are as indicated below:

Step 1: Initial denaturation at 94°C for 5 mins; Step 2: Denaturation at 94°C for 30 s; Step 3: Annealing at 56°C for 30 s; Step 4: Extension at 68°C for 1 min; Step 5: Final extension at 68°C for 5 mins. Steps 2-4 were repeated for a total of 35 cycles. Step 6: Hold at 12°C. The PCR products were resolved on a 1% (w/v) ethidium bromide-stained agarose gel run at 100 V. Gels were visualized via the Spectroline UV transilluminator SelectTM Series, and images were obtained via the AlphaImager 2200. Additional details for the RT‒PCR analyses are indicated in the respective figure legends, as appropriate.

### Growth phenotype analysis

Seedlings: Seeds were surface sterilized using 10% bleach three times for two minutes each, followed by three washes with sterile water, after which they were resuspended in 0.1% phytoagar and stratified in the dark at 4°C for at least two days. Seeds were then plated onto 1x MS agar plates (100 mm × 100 mm square plates; Fisher Scientific; Cat#FB0875711A), composed of 4.4 g/L Murashige and Skoog Basal Medium with vitamins (PhytoTech; M519), 0.5% sucrose (Sigma; S7903), 0.5 g/L MES (Sigma; M8250), and 0.7% PhytoAgar (PlantMedia; Cat#40100072-2), pH 5.7, and the plates were sealed with parafilm and incubated vertically in a reach-in growth chamber (Conviron Adaptis 1000, Canada) at 25 ± 2°C, 75% RH, 16 h light/8 h dark, and 100 µmoles/m2/s light intensity for two weeks. A separate set of seedlings was used to determine whole plant fresh and dry weight. A second and separate set of seedlings was used for shoot and root biomass analysis. For dry weight analysis, samples were freeze-dried in a lyophilizer (Labconco) overnight.

Adult plants: Stratified seeds were sown in peat pellets (Jiffy-7, Jiffy Products Ltd., Shippagan, Canada), 4–5 seeds per pellet, in trays, and covered with a translucent plastic dome to maintain high humidity. These were transferred to growth chambers under controlled conditions at 25 ± 2°C, 75% RH, 9 h light/15 h dark, and 100 µmoles/m2/s light intensity for the next 6 weeks. One week after sowing the seeds, the domes were removed, and the seedlings were thinned out leaving 1–2 seedlings per pellet. A second thinning out was performed at the end of week two, leaving 1 seedling per pellet through the end of the experiment. The plants were watered with Hoagland’s solution three times weekly for four weeks, and with water for the remainder of the experiment. Fresh and dry weights were determined after 6 weeks. For dry weight analysis, samples were dried in an oven at 55 ᴼC overnight. To characterize mutants for seed weight and number of seeds per silique, experiment was performed similarly under long-day conditions in a climate-controlled growth chamber for 9 weeks.

### Plant gene expression analysis (RT‒qPCR)

Harvested tissues (roots/shoots/leaves) were frozen in dry ice. RNA was isolated via the use of TRIzol® Reagent (Fisher Scientific; Cat# 15596018.) and quantified with a NanoDrop-ND1000 spectrophotometer (Thermo Fisher Scientific, Inc.). Two micrograms of total RNA was used for 1st strand cDNA synthesis after DNaseI treatment. The 2 µg RNA was incubated together with 2 µL of random decamers and an appropriate volume of DEPC-treated water to a final volume of 15 µL at 70°C for 5 minutes and then immediately transferred to ice. The cDNA synthesis mixture containing 10 mM dNTPs (2 µL), Promega RNase inhibitor (1 µL), Promega M-MLV-RT (1 µL), 5x Promega M-MLV Buffer (5 µL) and DEPC-treated water (1 µL) was subsequently added to a final volume of 25 µL, which was subsequently incubated at room temperature for 1 minute and then at 37°C for 60 minutes. For the qPCRs, 1 µL of cDNA was used in a total reaction volume of 20 µL containing 10 µL of SyBr Green mix, 2 µL of 5 µM primers (forward and reverse), and 7 µL of DEPC-treated water. There were two technical replicates per biological replicate. The reactions were performed using the ABI 7500 and 7500 Fast Real-Time PCR system v2.3

### Root exudate collection assays

For root exudate screening, root exudates were collected via a previously published method with modifications [36]. Briefly, Arabidopsis seedlings were grown initially in 1x MS medium (i.e., full-strength) in 12-well plates (USA Scientific; Cat# CC7682-7512) containing 0.5% sucrose for 12 days by placing ca. 5 seeds per well on an autoclaved mesh disc (McMaster-Carr; Cat# 1100t41) sitting on top of the medium, and the medium was changed to 0.5x MS medium (i.e., half-strength) containing no sucrose for 3 days, with the roots separated from the shoots by the autoclaved mesh. The root exudates were then collected and filter-sterilized through a 0.22 µm filter for further downstream processing. Throughout the experiment, the plates were sealed with parafilm and incubated in a growth chamber (Conviron Adaptis 1000, Canada) at 25 ± 2°C, 75% RH, 16 h light/8 h dark, and 100 µmoles/m2/s light intensity.

### Colorimetric/fluorometric quantitation of AAs

The total AAs content of the root exudates was assayed via the L-Amino Acid Quantitation Colorimetric/Fluorometric Kit (BioVision, Catalogue #K639-100) following the manufacturer’s instructions with modifications. A 12.5 μL reaction mixture containing 11.5 μL of L-amino acid assay buffer, 0.5 μL of L-amino acid probe, and 0.5 μL of L-amino acid enzyme mixture was added to each well containing 12.5 µL of the respective test samples or L-amino acid standard (for a total of 25 µL of reaction volume) for L-amino acid standard curve generation. The reactions were incubated in a SpectraMax® i3x (Molecular Devices) for 30 minutes at 37°C and assayed via the fluorometric method (Ex/Em = 535/587 nm). In our analysis, background correction was performed by subtracting the mean AA concentration for the unplanted samples from the concentration value for each experimental sample (i.e., wild-type and mutant root exudate samples). Occasionally, when this background subtraction yielded a negative value (i.e., below background), the AA concentration was set to zero and included in the final analysis.

### Liquid chromatography–mass spectrometry (LC–MS) analysis of AAs

The samples were vacuum-dried and reconstituted in 100 µL of buffer containing 0.1% formic acid (FA) and subsequently analysed on a Thermo Exploris 480 mass spectrometer via the ZipChip Interface. Standard curves of 20 AAs were generated to obtain absolute quantification of the concentration of AAs. For the root exudate samples, glycine data were omitted from the final analysis as MS media (Cat#M519, *Phyto*Technology Laboratories, LLC) for collecting root exudates contained background amounts of glycine (2 mg/L). For data analysis, raw data files for both the standard AAs and experimental samples were uploaded into Thermo XCalibur software (https://www.thermofisher.com/order/catalog/product/OPTON-30965#/OPTON-30965), and targeted peak detection was performed via the ICIS peak integration algorithm. Thermo quantitative analysis software (Quan browser) was then used to generate calibration curves, followed by determination of the concentration of the AAs in the root exudate and unplanted samples. LC‒MS analysis was performed at the W.M. Keck Biomedical Mass Spectrometry Laboratory at the University of Virginia.

### Amino acid uptake and secretion in planta

Arabidopsis seedlings were grown for two weeks via the root exudate collection assay described above in full-strength Murashige and Skoog (MS) media supplemented with 0.5% sucrose. The roots and shoots were separated, and each sample was transferred into 1 mL of full-strength MS media supplemented with 0.5% sucrose in 12-well plates for 5 h under agitation on a rocking surface (8 rpm). AA uptake was performed for 20 min in a mixture containing 1 mL of full-strength MS media supplemented with 0.5% sucrose and 100 µM glutamine + [3H]Gln, with a final specific activity of 3.7 kBq µmol–1. Efflux was performed for 20 min. Radioactivity in the root and shoot samples was then determined via a 1450 Microbeta Trilux Liquid Scintillation and Luminescence Counter (Perkin- ElmerTM Life Sciences).

### Amino acid extraction from root and leaf tissues for LC‒MS analysis

The plant material was collected in a 2 mL Eppendorf tube (round bottom) and lyophilized overnight in a lyophilizer (LABCONCO). The defined biomass was then weighed into a new Eppendorf tube, frozen in liquid nitrogen and homogenized with a tissue lyser (TissueLyser II, Qiagen) for 3 minutes at 15/s. Next, 700 µL of 100% methanol was added to each sample, and the mixture was vortexed briefly. The samples were subsequently shaken for 15 minutes at 4°C, and the tubes were opened approximately 2 minutes after the start of incubation to release the pressure in the tubes. Centrifugation was carried out at approximately 14000 rpm for 10 minutes, and the supernatant was transferred to a new tube. Next, 375 µL of chloroform was added to each supernatant, plus 750 µL of water (Nanopure). The samples were vortexed for 15 s each and then centrifuged for 15 min at 4000 rpm. The entire upper phase (polar phase) was collected into new Eppendorf tubes and then dried via a Savant DNA120 SpeedVac concentrator (Thermo Electron Corporation). These extracts were stored at -80°C until LC‒MS analysis.

### Beneficial rhizobacterium and growth conditions

*Pseudomonas simiae* WCS417r (previously known as *Pseudomonas fluorescens* WCS417r), which was initially isolated from lesions of wheat (*Triticum sativum*) roots [37], was maintained on LB plates supplemented with 50 µg ml−1 rifampicin. To prepare for the root inoculation experiments, a single colony was randomly picked from the appropriate plate and grown overnight in approximately 100 mL of LB at 28°C with shaking at 230 rpm until the cultures reached OD600 = 0.4–0.8. The cell cultures were harvested, washed three times in sterile water, and then adjusted to the required inoculation titre with sterile water.

### *Ps* WCS417r growth in root exudates

*Ps* WCS417r cells from overnight culture (OD600 ̴ 0.8) were washed three times in sterile water and resuspended in sterile water to OD600=0.2. Ten (10) µL was added to 90 µL of the unplanted growth medium (unplanted) or to 90 µL wild-type root exudates or 90 µL mutant root exudates, to a final OD600=0.02. The test samples were filter-sterilized through a 0.22 μm-filter before use. Growth was measured over a given duration with intermittent shaking, in the microplate reader SpectraMax® i3x (Molecular Devices).

### *Ps* WCS417r-mediated plant growth assay

Sterilized and stratified seeds were plated directly on 3MM paper placed inside a square plate (100 mm × 100 mm square plates; Fisher Scientific; Cat#FB0875711A) and wetted with half-strength Hoagland’s solution (i.e., Hoagland-only) or full-strength Hoagland’s solution mixed with equal volume of *Ps* WCS417r (OD600=0.4), yielding half-strength Hoagland’s solution containing *Ps* WCS417r OD600=0.2 (i.e., Hoagland + *Ps* WCS417r). For each genotype, twelve (12) seeds were sown per plate for *n*=5 plates. The plates were sealed with parafilm and incubated vertically in a reach-in growth chamber (Conviron Adaptis 1000, Canada) at 25 ± 2°C, 75% RH, 16 h light/8 h dark, and 100 µmoles/m2/s light intensity. Twelve days after incubation, shoots or roots from one plate were pooled into one sample for shoot and root fresh weight determination.

### Statistical analysis

Data analyses were performed via JASP open-source software v 0.14 and Excel, and graphs were generated via GraphPad Prism and Excel. A two-sided Student’s *t* test was performed for statistical comparison of two means, or a Welch’s *t* test was used for comparison of two means with unequal variances, when relevant. For comparisons of more than two means, one-way ANOVA followed by Tukey’s post hoc test or a Kruskal‒ Wallis test for unequal variances followed by Dunn’s post hoc test was performed, as indicated in the relevant figure legends.

For statistical analysis of bacterial growth curves, the CGGC (Comparison of Groups of Growth Curves) permutation test [38] was used to compare pairs of samples (e.g., wildtype vs. *umamit30*) over the course of growth (e.g., 24 hours). The test statistic (mean *t*) is the two-sample *t* statistic used to compare the OD600 values between the two groups at each hour, averaged over the course of growth (e.g., 24 hours). A P value was obtained for the test statistic via simulation. The samples were randomly allocated to each of the two groups, and the mean *t* was recalculated for 10 000 datasets generated through this permutation. The P value is the proportion of permutations where the mean *t* is greater in absolute value than the mean *t* for the original dataset. (That is, the number of times the absolute mean *t* from the permutations is greater than the absolute mean *t* for the original dataset, all divided by 10 000.) This P value was interpreted as the probability of obtaining the mean *t* obtained for the original dataset if the null hypothesis was true.

## Results

### *UMAMIT30* is expressed in roots and shoots under steady-state conditions

To identify AA exporters expressed in Arabidopsis roots and potentially contributing plant-derived AAs in the rhizosphere, we analysed publicly available microarray datasets [39, 40, 41] to examine the expression profiles of several *UMAMIT* genes across different tissues and developmental stages. This examination revealed that *UMAMIT30* is strongly expressed in Arabidopsis roots relative to its expression in other tissues during the seedling and vegetative stages (**Figure S1A**). To confirm these expression patterns, we performed RT‒qPCR on RNA derived from the roots and shoots/leaves of wild-type seedlings and adult plants growing under standard conditions (i.e., not subjected to any experimental treatment). These analyses revealed very strong expression of the *UMAMIT30* gene in the root tissues compared with the expression in the leaves (**Figure 1A and 1B**). Notably, the publicly available root tissue-specific expression data revealed that *UMAMIT30* expression is strongest within the vasculature (**Figure S1B**). This expression pattern is consistent with a potential role for UMAMIT30 in unloading AAs from the phloem into the stele apoplast, an important step in the transport of AAs from root tissues to the surrounding medium. More recent studies using a single cell RNA sequencing approach to assess gene expression show that UMAMIT30 mRNA is detected in phloem parenchyma cells of the leaf (**Figure 1SC**) and in pericycle and phloem cells of the root (**Figure 1SD**) [42, 43, 44].

**Figure 1:**
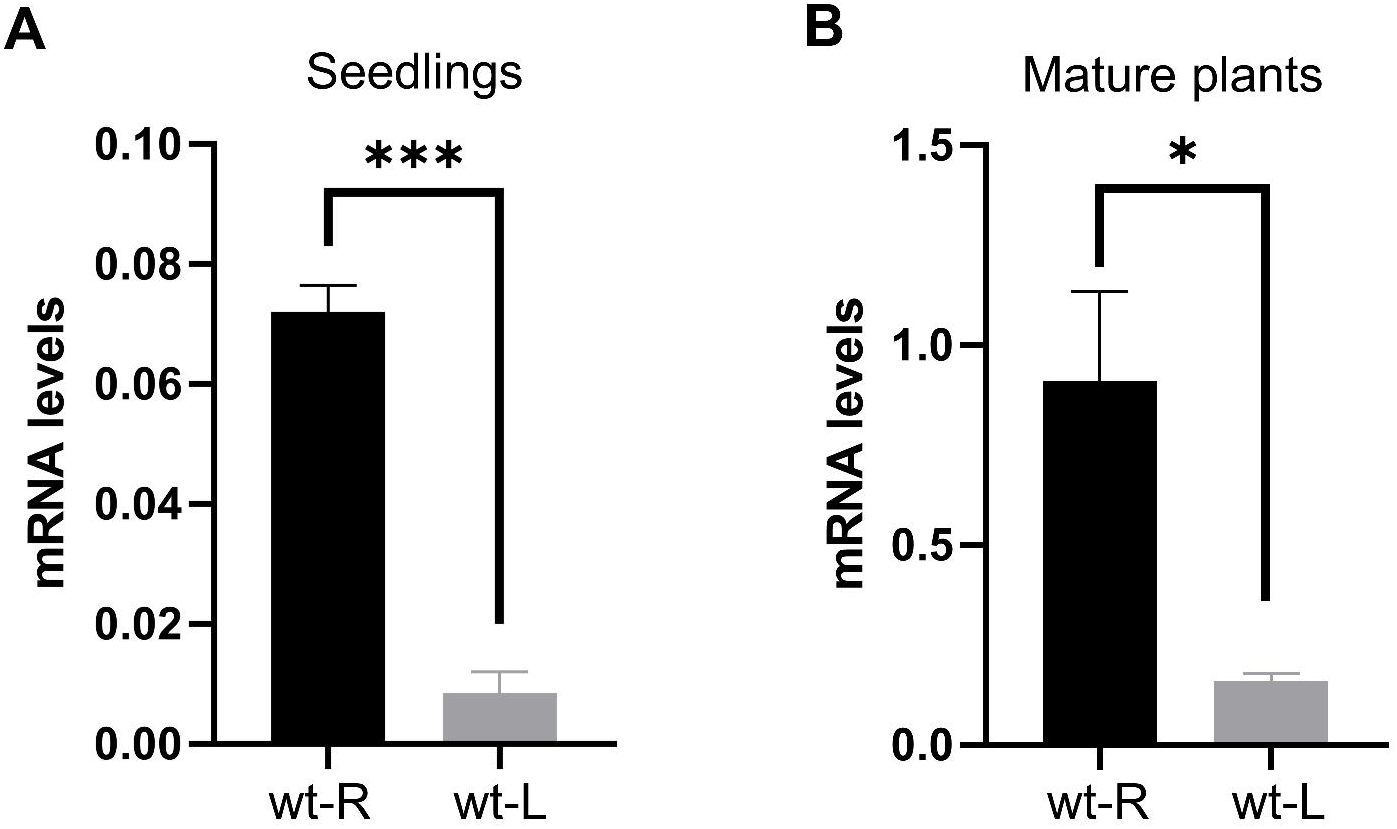
*UMAMIT30* gene expression in wild-type roots (wt-R) and leaves (wt-L). RT-qPCR data obtained from 14-day-old wild-type seedlings **(A)** and 6-week-old wild-type plants (i.e., mature plants) **(B)**. *UMAMIT30* gene expression levels were normalized against *ACTIN2*. AVR ± SE (*n*=3). Statistically significant differences (*p* ≤ 0.001) determined by a two-sided Student’s *t*-test, or *p* ≤ 0.05 by Welch’s *t*-test, are indicated by (***) and (*), respectively. Similar results were obtained in three independent experiments.

### *UMAMIT30* expression knockouts

To analyse *UMAMIT30* function in planta, two homozygous Arabidopsis T-DNA insertion lines (SALK_140547C and SALK_146977C, hereafter referred to as *umamit30-1* and *umamit30-2*) were identified by PCR genotyping. To test for potential impact of these T-DNA insertions on UMAMIT30, gene expression analysis was carried out via PCR using *UMAMIT30* specific primers **(Table S1)**. Retro-transcription of RNA samples followed by polymerase chain reaction (RT-PCR) analysis showed that *UMAMIT30* expression was knocked out in both *umamit30* mutants, as *UMAMIT30*-specific PCR products were not detected in either of the insertional lines whereas the *UMAMIT30*-specific PCR product was present in PCR reactions using cDNA obtained from wild-type plants (**Figure 2A and 2B**).

**Figure 2:**
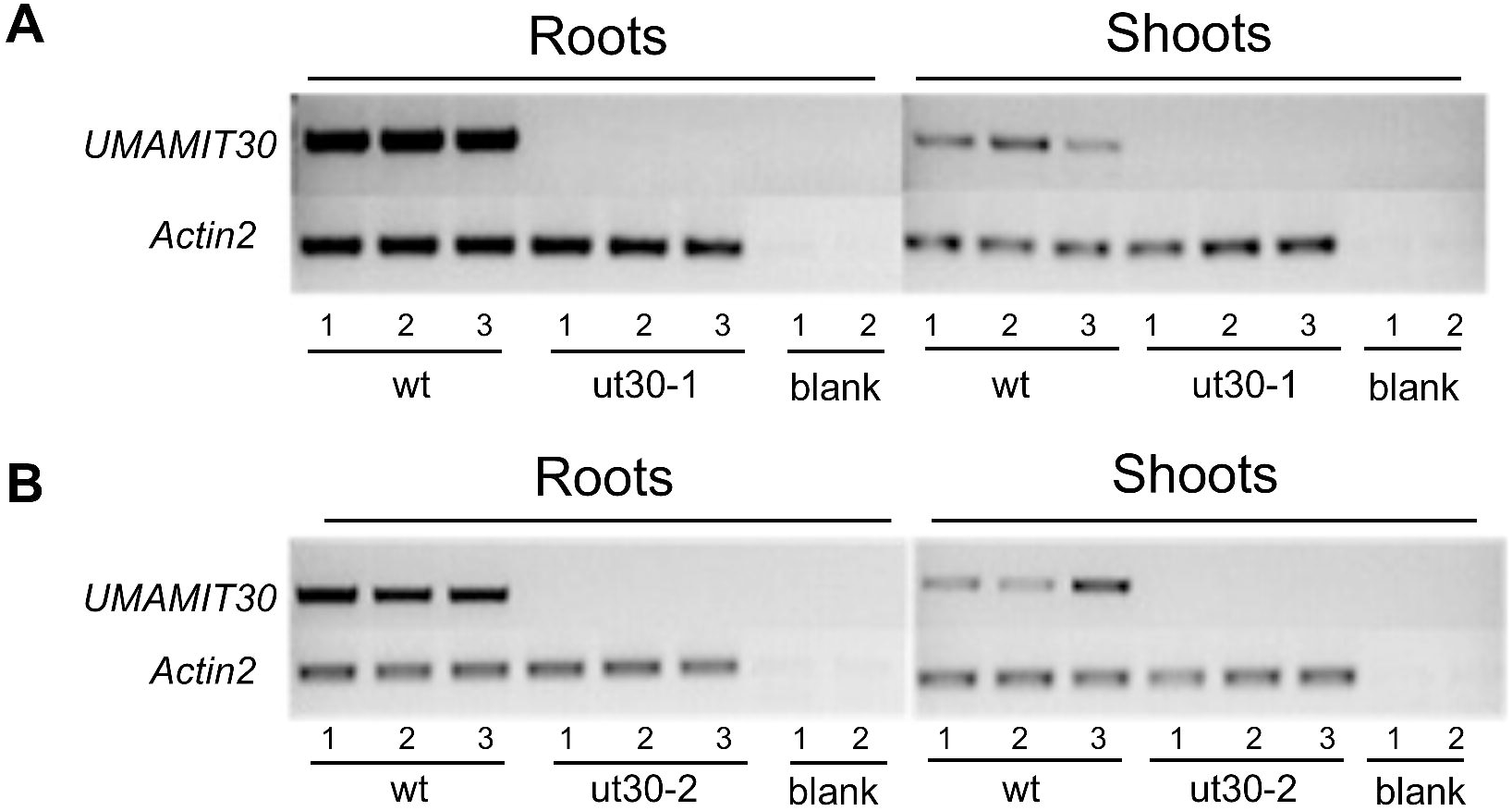
***UMAMIT30* gene expression in wild-type and *umamit30* plants.** RT-PCR analysis of *UMAMIT30* gene expression in roots and shoots of wild-type and **(A)** ut30-1 (*umamit30*-1; Salk_140547C) or **(B)** ut30-2 (*umamit30-2*; Salk_146977C) in 14-day-old seedlings. *ACTIN2* expression was used as a control for cDNA input. Blank = no template control. Corresponding uncropped full-length gel images are shown in Figure S2A and S2B.

### UMAMIT30 contributes to AA root exudation

Because the *UMAMIT30* gene is strongly expressed in the root vasculature (**Figure S1B**), we hypothesized that the protein functions in root-to-medium AA transport. A root exudate collection assay, adapted from a previous study [36] and similar to the one used by Besnard et al. [29] to assess UMAMIT14 and UMAMIT18 AA secretion in Arabidopsis, was used to test this hypothesis (see Materials and Methods). While root biomass remained unchanged in both *umamit30* insertional mutants (**Figure 3A and 3B**), analysis of the root exudates revealed a significant depletion of AAs (**Figure 3C and 3D**). These data potentially indicate that UMAMIT30 may function in exporting AA from Arabidopsis roots. To further confirm the reliability of the assay, the impaired root AAs secretion phenotype of *umamit14* plants [29] was used as a control (**Figure S3A**). In addition, the AA concentrations in the root exudates of *umamit05* suggest that UMAMIT05, which is capable of exporting a single AA when expressed in yeast cells [32], may not contribute to the secretion of AAs into the root exudate (**Figure S3B**). The analyses of *UMAMIT14* and *UMAMIT05* expression revealed that these genes were not expressed in the roots of the respective T-DNA insertional lines (**Figure S3C and S3D**). Taken together, these results confirm the root AAs export function of UMAMIT14 and reveal the lack of root exudate phenotype of UMAMIT05. Overall, these data show the reliability of the assay in detecting differences in AA concentrations in the root exudates of wild-type and mutant plants.

**Figure 3:**
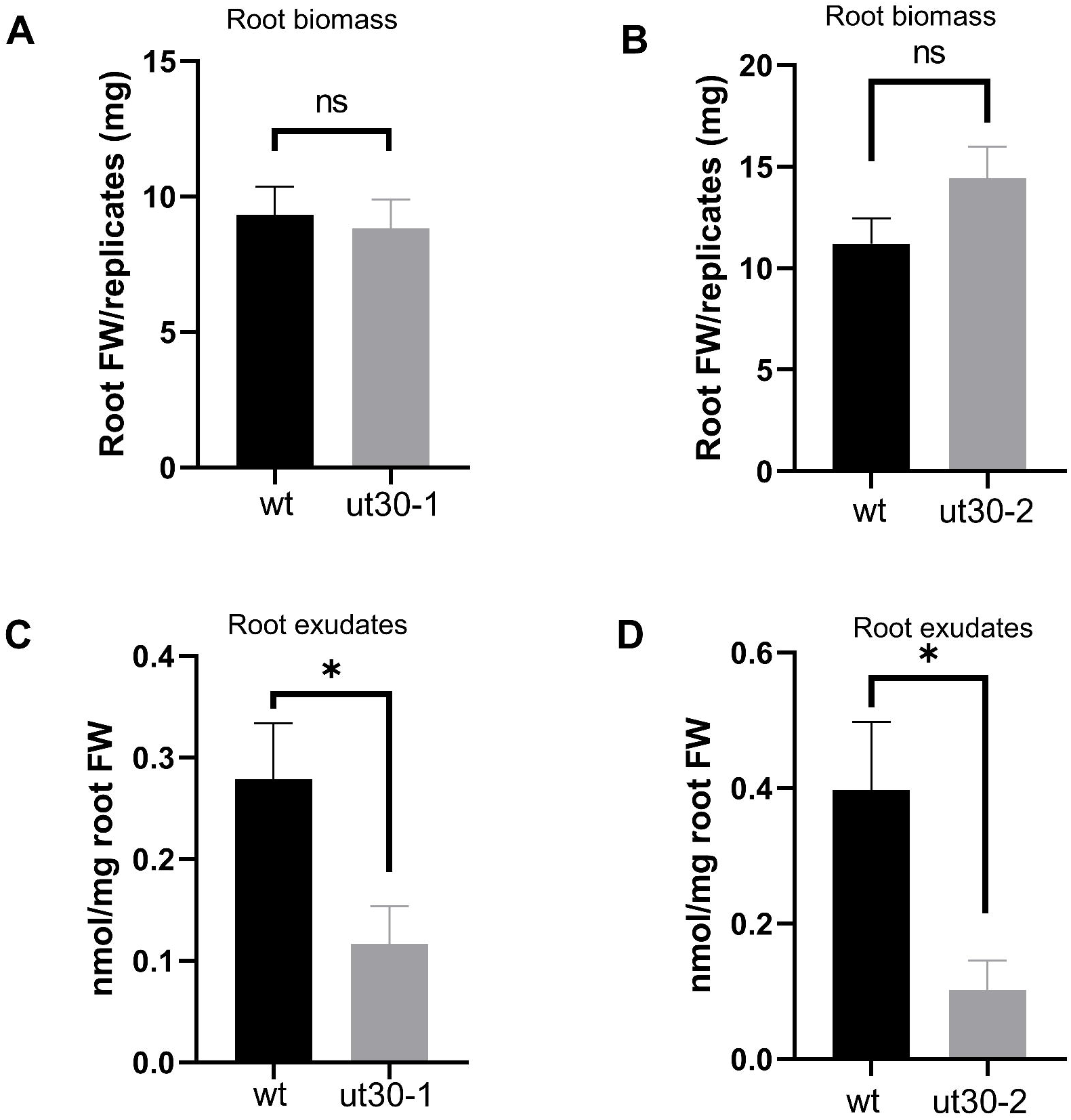
Root biomass (A, B) and concentration of total free amino acids in root exudates (C, **D)** of wild-type, ut30-1 (*umamit30*-1; Salk_140547C) and ut30-2 (*umamit30-2*; Salk_146977C) in 14-day-old seedlings. AVR ± SE (*n*=6). Statistically significant differences (*p* ≤ 0.05) between wt and ut30 determined by a two-sided Student’s *t*-test are indicated by (*). ns: Non-statistically significant difference. Similar results were obtained in three of three independent experiments.

Arabidopsis AA transporters have a relatively broad specificity for substrates. Thus, to assess the AA accumulation signatures of *umamit30* root exudates, filter-sterilized root exudates from both the wildtype and *umamit30* were further analysed via liquid chromatography coupled with mass spectrometry (LC‒MS). Consistent with previous studies showing that most UMAMIT transporters have low substrate specificity [32], the AA profile of the *umamit30* root exudates showed a significant reduction in the concentration of various AAs relative to the wildtype levels (**Figure 4**). In particular, the concentrations of asparagine and glutamine decreased significantly and likely contributed the most to the decline in total AAs levels (Figure 3C–D).

**Figure 4:**
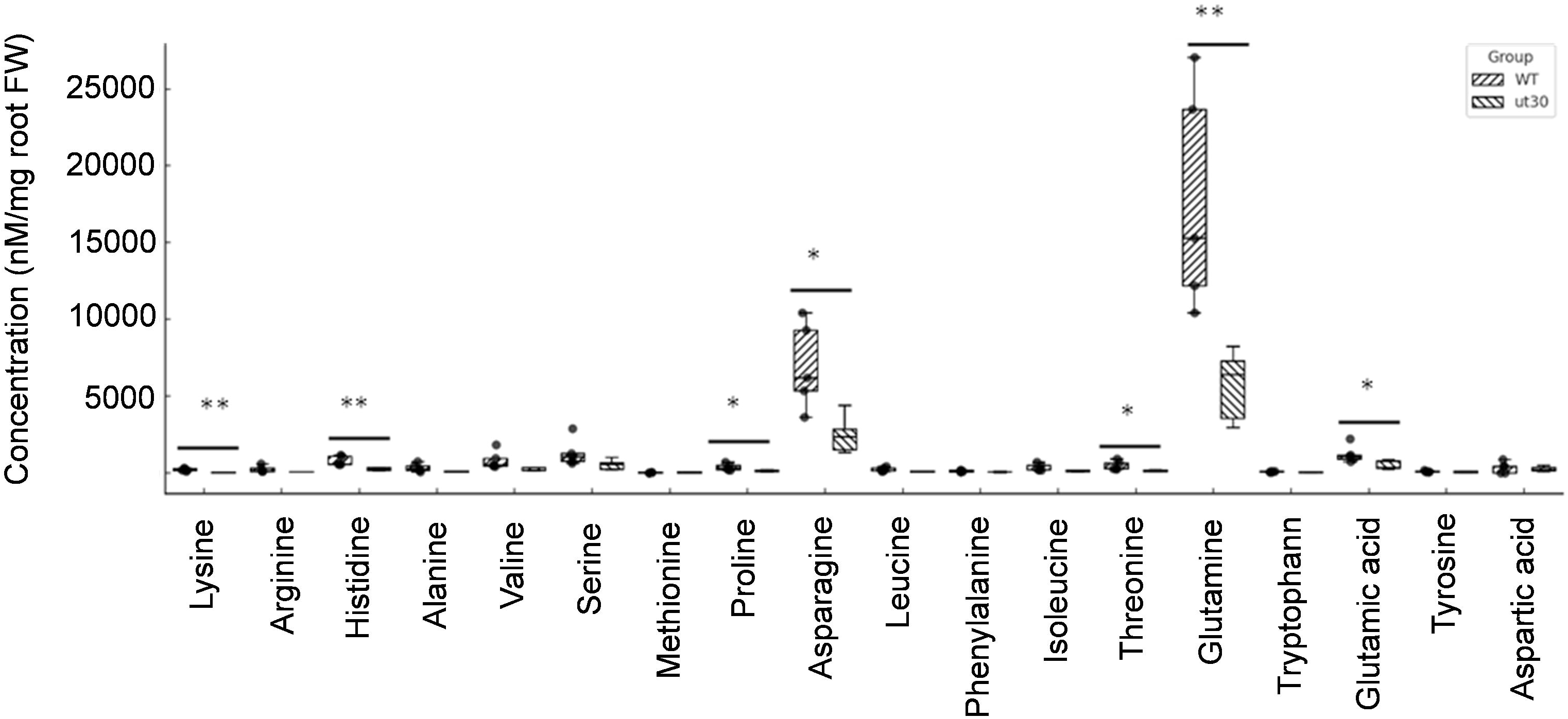
Concentrations of individual amino acids in root exudates. Wild-type (WT) and *umamit30-1* (*ut30*; Salk_140547C) liquid root exudates were analyzed via Liquid-Chromatography coupled to Mass Spectrometry. Concentrations (nmol/mg) were normalized by root fresh weight (FW). Statistically significant differences between WT and ut30 were determined using a two-sided Student’s *t*-test (*n*=5) and are indicated by (*) for *p* ≤ 0.05 and (**) for *p* ≤ 0.01. Two of two experiments showed similar results.

### Loss of UMAMIT30 function does not impact plant growth

Mutations in AA transporters may alter AA homeostasis and impact growth phenotypes [45, 46]. To examine this possibility, we visually inspected *umamit30* plants and found no apparent growth defects in either roots or shoots (**Figure S4A**). Furthermore, biomass analyses revealed no significant differences in whole seedling fresh or dry weight (**Figure S4B; Figure S4C**). Similarly, there was no significant difference in shoot fresh or dry weight (**Figure S4D**; **Figure S4E**) or root fresh weight (**Figure S4F**) under *in vitro* growth conditions. A similar analysis of the shoots of mature plants grown in peat pellets revealed that the fresh (**Figure S5A and C**) and dry weights (**Figure S5B and D**) of the *umamit30* plants were similar to those of the wild-type plants. In addition, the phenotypic characterization of 9-week-old plants grown under long-day conditions in a climate-controlled growth chamber revealed no differences in plant height, biomass, seed weight or number of seeds per silique between the mutants and the wild-type plants (**Table 1**). Thus, the depletion of AAs in root exudates in *umamit30* is likely not explained by altered AAs homeostasis, which would have led to altered growth phenotypes.

**Table 1:** Morphological characterization of wild-type and *umamit30* plants Plant height, biomass, seed weight and seed number per silique were recorded nine-weeks post germination under long day conditions. One-way ANOVA in conjunction with Tukey’s post-hoc test was used for height, biomass, and number of seeds/silique. Seed weight (weight of 100 seeds) data was analyzed using Kruskal—Wallis test followed by Dunn’s post-hoc test. AVR ± S.E. (*n*=4). Same letter (a), across all genotypes, denotes no statistically significant differences between genotypes (*p* ≤ 0.05).

|  | Plant height<br>(cm) | Biomass, FW<br>(g) | Weight of 100<br>seeds (mg) | Number of<br>seeds per<br>silique |
| --- | --- | --- | --- | --- |
| **Wild-type** | 53.5 ± 0.65<br>(a) | 3.7 ± 0.15<br>(a) | 2.1 ± 0.17<br>(a) | 44.0 ± 2.68<br>(a) |
| ***umamit30-1*** | 52.0 ± 1.22<br>(a) | 3.3 ± 0.30<br>(a) | 2.4 ± 0.03<br>(a) | 43.0 ± 2.16<br>(a) |
| ***umamit30-2*** | 50.3 ± 2.06<br>(a) | 3.5 ± 0.21<br>(a) | 2.4 ± 0.05<br>(a) | 42.3 ± 1.25<br>(a) |

### UMAMIT30 contributes to uploading AAs to shoots

As *UMAMIT30* is strongly expressed in the root tissues where it may export AAs out of the root vasculature, we expected that AAs may accumulate in roots in the absence of *UMAMIT30* function. To test this hypothesis, we quantified the total concentration of AAs in the root and shoot tissues of both wild-type and *umamit30-1* seedlings. Surprisingly, no significant difference in the AAs content between the roots of wild-type and *umamit30* seedlings was detected (**Figure 5A**). Instead, the shoot AA content was elevated in *umamit30* seedlings (**Figure 5B**). These data suggest that UMAMIT30 may play an important role in secreting AAs into the apoplast from where they can be uploaded to the phloem by companion cells localized active transporters.

**Figure 5:**
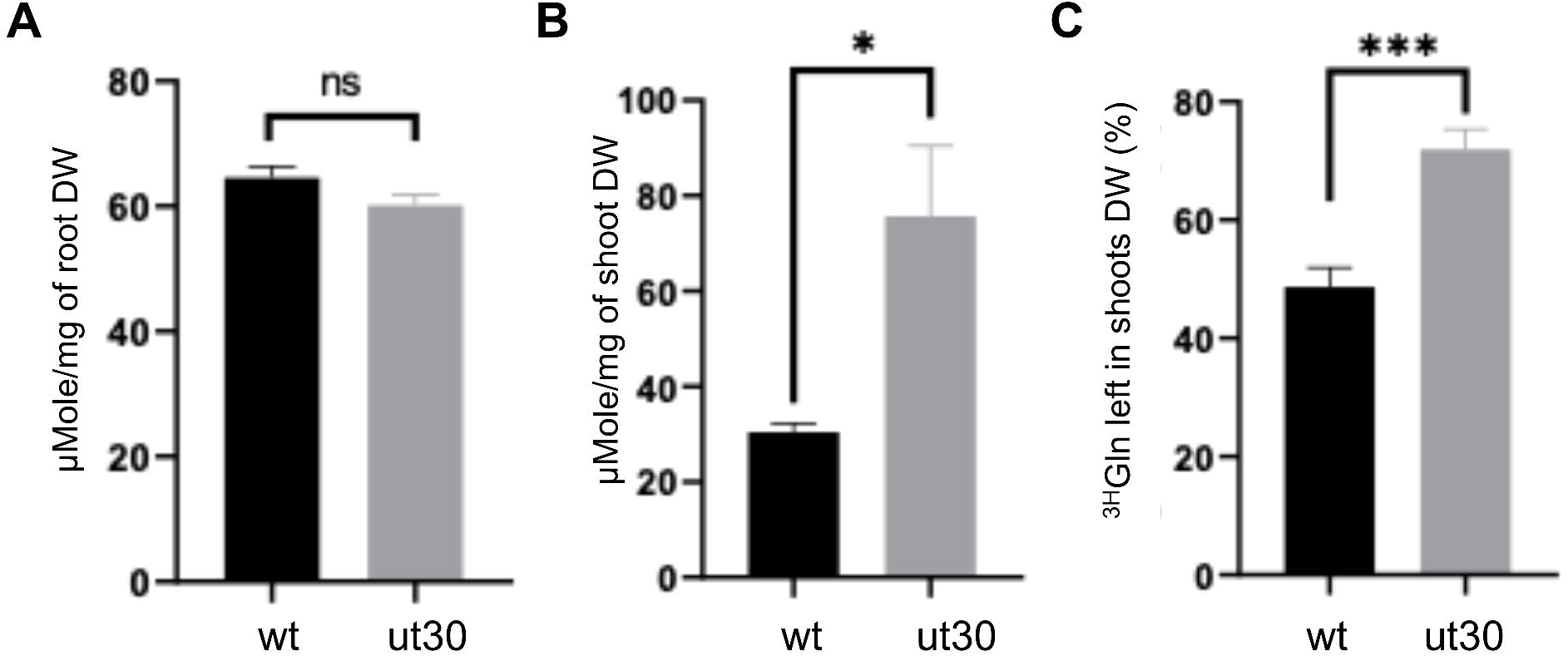
R**e**tention **of amino acids in *umamit30* shoots.** Concentration of total free amino acids in roots **(A)** and shoots **(B)** of wild-type and *umamit30-1* (ut30) were quantified using a fluorometric assay (BioVision total L-AAs quantification) as described in Material and Methods. AVR ± SE (*n*=3). Statistically significant difference between means (*p ≤* 0.05) indicated by (*); two-sided Students *t*-test. ns: Non-statistically significant difference. **(C)** Gln secretion was assessed as a percentage of uptake and informed as ^3H^Gln left in samples. Higher signal indicates lower Gln secretion. AVR ± SE (*n*=5). Statistically significant difference (*p* ≤ 0.001) is indicated by (***); two-sided Students *t*-test. Two of two independent experiments showed similar results.

To gain understanding on UMAMIT30 contributions to AAs transport, the efflux of radiolabelled glutamine, i.e., [3H]Gln, was assessed in wild-type and *umamit30* seedlings. While root efflux analysis was inconclusive, [3H]Gln efflux from shoots decreased significantly in the *umamit30-1* seedlings (**Figure 5C**). These data, together with data showing that *umamit30-1* seedlings accumulate more AAs in their shoots than wild-type seedlings, suggest that the depletion of AAs in the root exudates of *umamit30* seedlings may be related to impaired leaf phloem uploading. In light of the AAs shoot accumulation phenotype of *umamit30* seedlings, the potential contribution of UMAMIT30 to root exudation could not be determined conclusively in this study and would require tissue-specific suppression and induction of UMAMIT30 expression.

### AA depletion in *umamit30* root exudates does not affect root―*Ps* WCS417r interactions

A few examples of the functional significance of plant AA transporters in mediating plant―biotic interactions are known. For instance, the AA transporter CAT6 and some transporters belonging to the AAP family are known to be transcriptionally induced during nematode invasion of plant tissues [47, 48, 49]. In Arabidopsis *aap3* and *aap6* mutants, nematode root infestations are reduced compared with that in wild-type plants [49, 50, 51]. To determine whether the decreased AA content in root exudates of *umamit30* could hinder beneficial microbe interactions, the growth of *Ps* WCS417r in wild-type and *umamit30* root exudates was examined. The results revealed that *umamit30* root exudates (**Figure S6A**), like *umamit14* root exudates included here as a control **(Figure S6B)**, were just as efficient as wild-type root exudates in supporting *Ps* WCS417r growth under *in vitro* conditions. By way of comparison, the AAs-rich exudates of *lht1* supported significantly more bacterial growth than root exudates obtained from*umamit30*, *umamit14*, and wild-type seedlings **(Figure S6B)**.

To further examine whether decreased exudation of AAs could impact beneficial bacteria-mediated plant growth, the roots of both the *umamit30* and *umamit14* mutants, along with the wildtype, were inoculated with *Ps* WCS417r. The data showed that *Ps* WCS417r- mediated plant growth promotion was not impaired in either mutant (**Figure 6**). These results indicate that lower glutamine exudation, at least under the conditions tested, does not compromise *Ps* WCS417r-mediated plant growth promotion.

**Figure 6:**
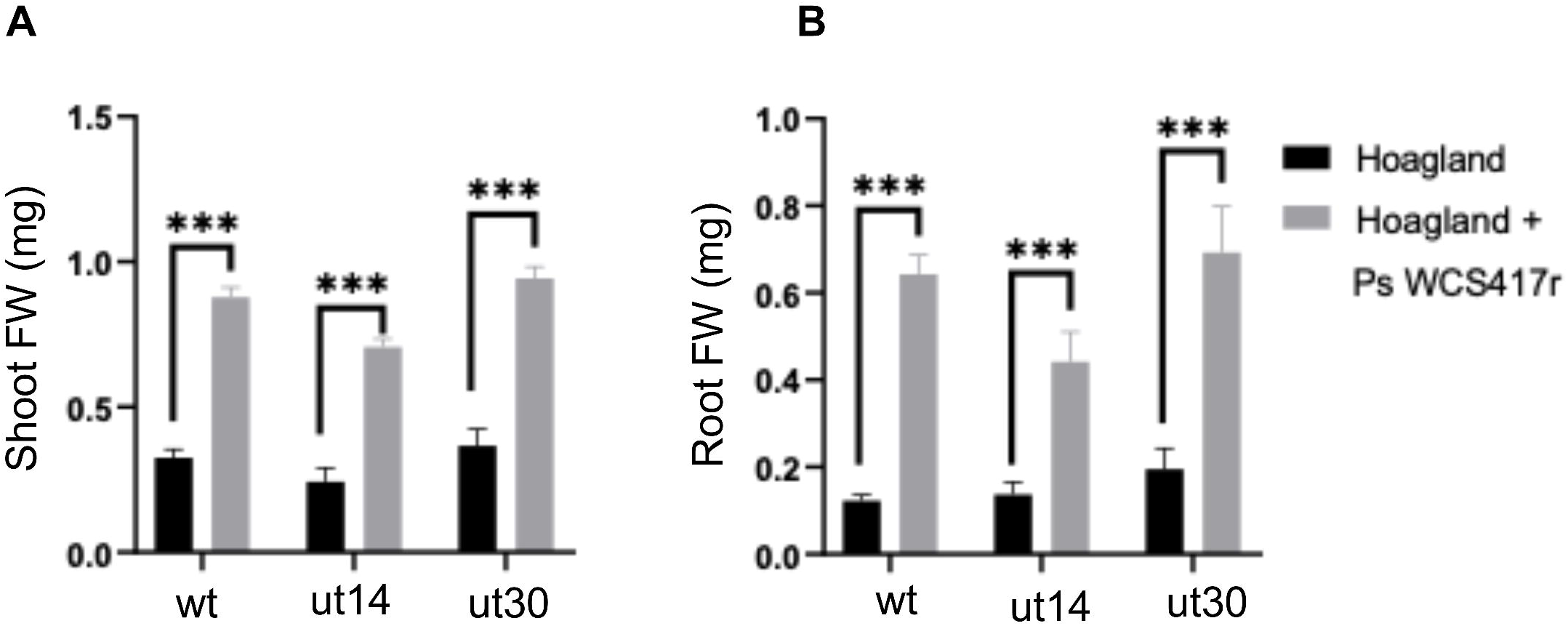
P***s* WCS417r-mediated plant growth promotion.** Arabidopsis shoot **(A)** and root **(B)** fresh weight (FW) recorded in 12-day old wild-type, *umamit14,* and *umamit30* seedlings growing on filter paper substrate supplemented with Hoagland-only or Hoagland + *Ps* WCS417r. AVR ± SE (*n*=5) of 12 seedlings per sample pooled together. Statistically significant differences (*p* ≤ 0.001) are indicated by (***); two-sided Student’s *t*-test. Two of two independent experiment showed similar results.

## Discussion

### UMAMIT30 contributes to AA root exudation by uploading AAs to the leaf phloem

In this study, we identify *UMAMIT30* as a contributor to AAs exudation in Arabidopsis roots, expanding our understanding of the UMAMIT family of AA transporters in shaping rhizosphere nutrient composition. Our findings show that *UMAMIT30* is highly expressed in Arabidopsis leaves and roots (**Figure 1 and 2**). In addition, publicly available single cell RNA sequencing data show that *UMAMIT30* is expressed in phloem parenchymal cells of Arabidopsis leaves and roots (**Figure S1C–D**). Since AAs are mostly synthesized in mesophyll cells of mature leaves, in order for AAs to be exuded by roots they must first be uploaded to the leaf phloem. In Arabidopsis, phloem parenchymal cells and phloem companion cells are not connected symplastically via plasmodesmata. Thus, AAs synthesized in mesophyll cells must first leak out of phloem parenchyma cells and then be taken up by active transporters to load them into phloem companion cells and sieve elements [52, 53]. Since *UMAMIT30* is expressed in phloem parenchyma cells of leaves (**Figure S1C**), its loss of function promotes the accumulation of AAs in green tissues and compromises glutamine secretion (**Figure 5B–C**). Thus, the function of UMAMIT30 in leaves could be to allow AAs to leak out of phloem parenchyma cells from where they can be taken up into companion cells by active transporters. A similar function has been proposed for facilitator transporters that move sugars out of parenchymal cells [42].

Like UMAMIT14 and UMAMIT18, which facilitate AAs efflux from the phloem into the root cortex and stele apoplast [29], UMAMIT30 could also contribute to secrete AAs into the surrounding medium. Like *UMAMIT14*, which localizes to pericycle and phloem companion cells, *UMAMIT30* shows strong vascular expression, suggesting its participation in the root exudation pathway, possibly in phloem unloading or redistribution of AAs toward the outer root cell layers. Whether or not UMAMIT30 also contributes to unloading AAs from the root vasculature to the endodermis, the cortex and the epidermis of the root remains unknown. Such hypothesis cannot be tested in the *umamit30* insertional mutant because phloem loading in leaves, a step upstream of phloem unloading in the root, is compromised in this mutant, hindering our ability to address this question. Addressing a potential role for UMAMIT30 in phloem unloading in the root will require the use of tissue-specific gene expression downregulation, which is out of the scope of the study presented here.

### Drop in concentration of several AAs, especially glutamine, does not impact the ability of *Ps* WCS417r to promote plant growth

The AA profile of *umamit30* root exudates revealed that the concentrations of several AAs, including lysine, histidine, proline, asparagine, threonine, glutamine, and glutamic acid were significantly reduced. (**Figure 4**). This broad specificity mirrors the substrate promiscuity observed in other UMAMITs, including UMAMIT14, UMAMIT18, and UMAMIT30 itself when heterologously expressed in yeast [32]. These patterns suggest that UMAMIT30 acts as a broad-range AA facilitator, and its loss of function alters the overall profile of AAs available in the rhizosphere. Despite the lower concentrations of AA in the exudates of *umamit30*, the growth of the *Ps* WCS417r under *in vitro* conditions (**Figure S6A**), or its ability to promote plant growth in Arabidopsis remains unchanged (**Figure 6**). This suggests that *Ps* WCS417r can tolerate or adapt to a range of AA concentrations in the rhizosphere, provided that the composition remains broadly supportive. In contrast, the *lht1* mutant, which overaccumulates glutamine in root exudates, supports increased bacterial growth *in vitro* (**Figure S6B**) but impairs plant growth promotion (25), suggesting that not just quantity but also AA composition critically shapes this root-bacteria interaction. Thus, while exudates with excess glutamine may compromise *Ps* WCS417r metabolism in its ability to promote plant growth, the modest, distributed lower concentrations of multiple AAs in *umamit30* exudates appears to maintain a favorable rhizosphere environment that promotes plant growth.

Our findings also reinforce the layered model of AA secretion in roots, as proposed in Besnard et al. [29], wherein multiple UMAMITs operate at distinct spatial checkpoints: UMAMIT30 in the vasculature for phloem unloading (**Figure 5B–C**), UMAMIT14 and UMAMIT18 in the root stele for radial transport, and other UMAMITs such as UMAMIT04, UMAMIT06, UMAMIT37, and UMAMIT42 in root epidermal cells and root hairs for final release into the rhizosphere. That *UMAMIT30* mutants show no growth defects (**Figure S4, Figure S5, Table 1**) suggests that its role in AA export may be functionally redundant or conditionally important, perhaps becoming critical under nutrient stress, specific developmental stages, or under biotic challenge.

Together, our data support a model in which UMAMIT30 facilitates the uploading of AAs from leaf apoplast, enabling their eventual secretion into the rhizosphere. Whether or not UAMIT30 also directly contributes to AAs root exudation is still unknown. This process shapes the nutrient landscape of the root-soil interface, with implications for microbial colonization and symbiosis. Importantly, our findings emphasize that the qualitative composition of exudates governs microbe-mediated outcomes, and that altering this balance can either impair or preserve beneficial plant-microbe interactions. Future work could take advantage of tissue-specific knockdowns or promoter swaps to dissect the precise step at which UMAMIT30 acts within the exudation pathway and explore how changes in AA profiles influence microbial gene expression, metabolism, and community dynamics. This would advance our understanding of how plants use transporter networks to engineer their rhizosphere and regulate microbial partnerships.

## Conclusions

In this study, we characterized the role of *UMAMIT30*, which encodes an AA transporter in Arabidopsis, in modulating the AAs composition of root exudates. Through a combination of expression analysis, radiolabelled AA transport assays, and bacterial growth assays, we demonstrated that *umamit30* loss of function mutants accumulate lower concentrations of AAs, especially glutamine, in root exudates. This phenotype suggests a defect in phloem loading. In spite of the altered amino acidic composition in root exudates, *Ps* WCS417r-mediated plant-growth promotion remained unaffected in *umamit30*. Taken together with data obtained from the *lht1* mutant [25], these findings indicate that while an excess of glutamine negatively impacts the ability of *Ps* WCS417r to promote plant growth, relatively low levels of glutamine do not have the opposite effect. Further studies will be necessary to establish if changes in the concentration of other AAs could improve plant growth promotion mediated by *Ps* WCS417r or other plant-growth promoting bacteria.

## List of abbreviations

Ps WCS417r: Pseudomonas simiae WCS417r
UMAMIT: <u>U</u>sually <u>M</u>ultiple <u>A</u>cids <u>M</u>ove In and out <u>T</u>ransporters
AA: amino acid
Aas: amino acids
LC-MS: Liquid chromatography–mass spectrometry

## Declarations

### Ethics approval and consent to participate

Not applicable

### Consent for publication

Not applicable

### Availability of data and material

The data sets used and/or analysed during the current study are available from the corresponding author upon reasonable request.

### Author Contributions

IDKA and CHD conceived and designed the experiments. IDKA performed experiments and analyzed data. PK assisted with uptake and secretion experiments using radiolabelled amino acids, data analysis, and data formatting. CHD assisted with gene expression experiments. IDKA wrote the original draft of the manuscript. CHD wrote the final form of the manuscript. All authors read and approved the final version for publication.

## Acknowledgements

We thank Dr. Nishikant Wase of the W.M. Keck Biomedical Mass Spectrometry Laboratory at the University of Virginia School of Medicine for LC-MS quantification of AA. The W.M. Keck Biomedical Mass Spectrometry Laboratory is funded by a grant from the University of Virginia School of Medicine.

---

**Figure S1:**
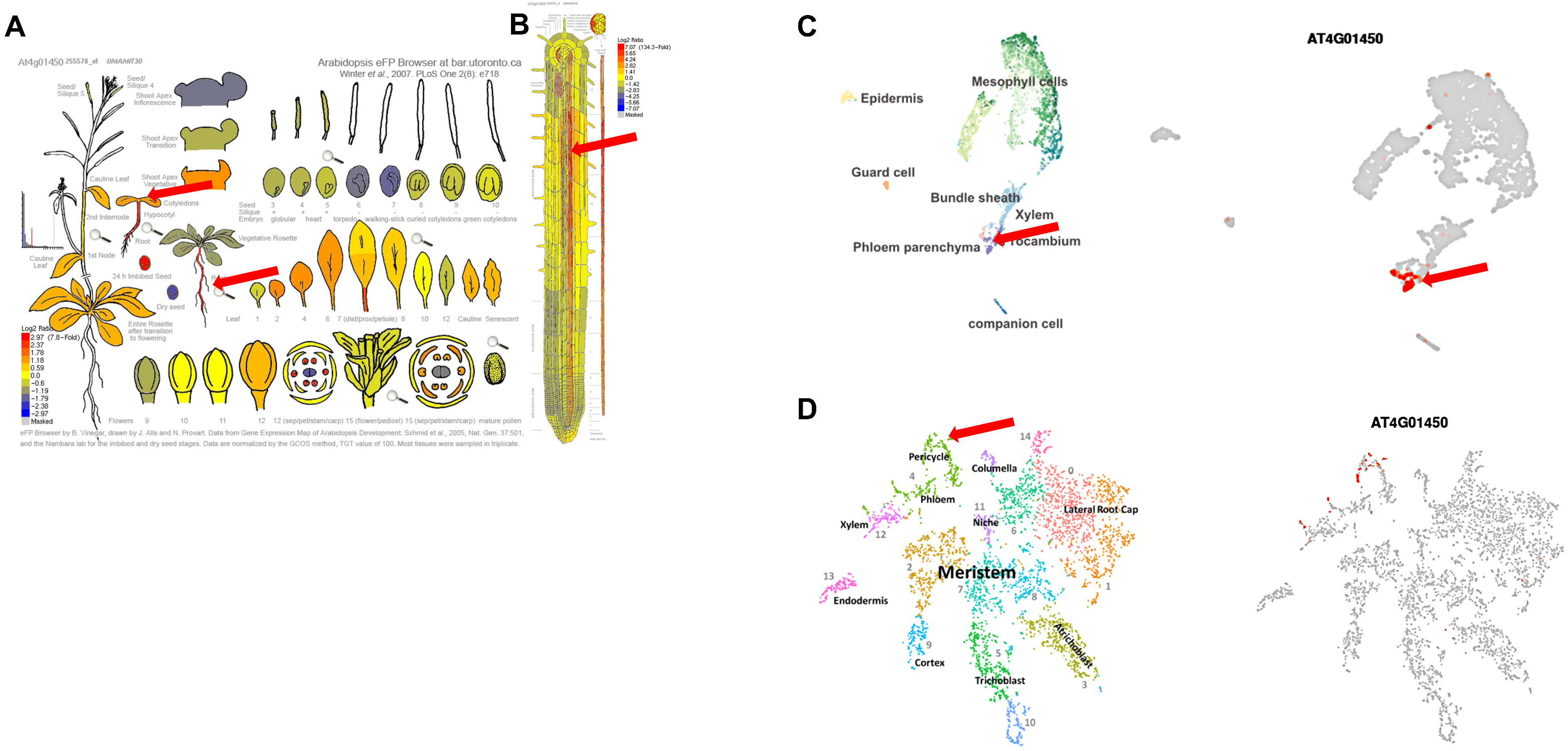
Tissue- and cell-type specific expression of *UMAMIT30*. Images of Arabidopsis Developmental Map **(A)** and Arabidopsis root tissue-specific gene expression pattern **(B)** generated by the Bio-Analytics Resource Center at the University of Toronto (http://bar.utoronto.ca/efp/cgi-bin/efpWeb.cgi) showing *UMAMIT30* (locus ID At4g01450) gene expression based on publicly-available microarray data sets [40]. Signals greater than 20 expression units were color-coded in relation to their average levels in the respective data sets [39, 41]. Images of single cell *UMAMIT30* gene expression in leaves **(C)** and roots **(D)** derived from single-cell RNA-sequencing data sets obtained by Kim et al. 2021 [42] and Denyer et al. 2019 [43], respectively. UMAP plots where generated by The Plant Single Cell RNA sequencing browser tool [44]. Red arrows point to tissues (A and B) and cell-types (C and D) where *UMAMIT30* expression was detected.

**Figure S2A:**
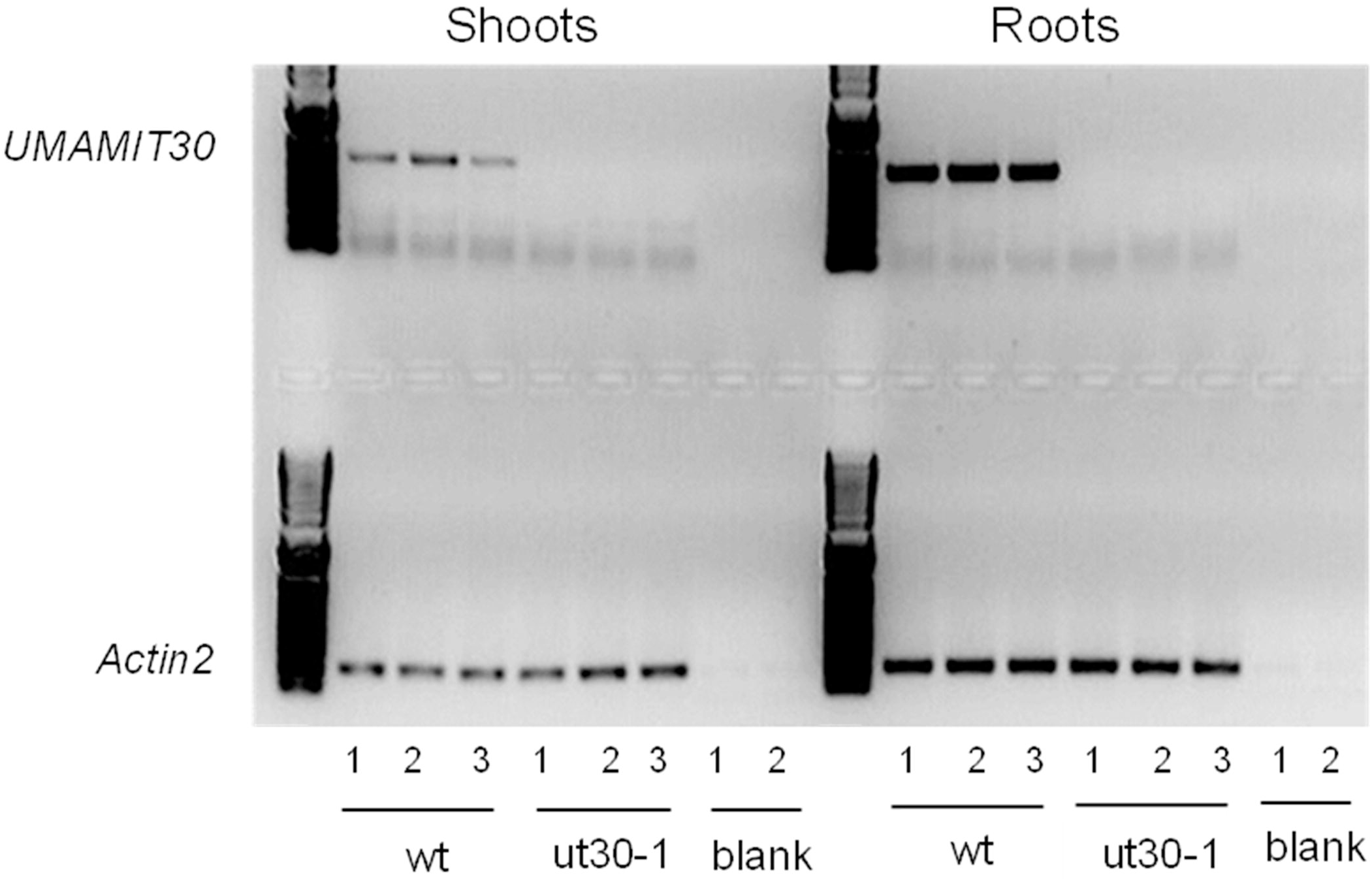
Analysis of *UMAMIT30* gene expression in wild-type and *umamit30-1* plants by RT-PCR. Blank = no template control. Source gel (uncropped full-length gel) for Figure 2A in the main text. RT-PCR primers are located in Table S1.

**Figure S2B:**
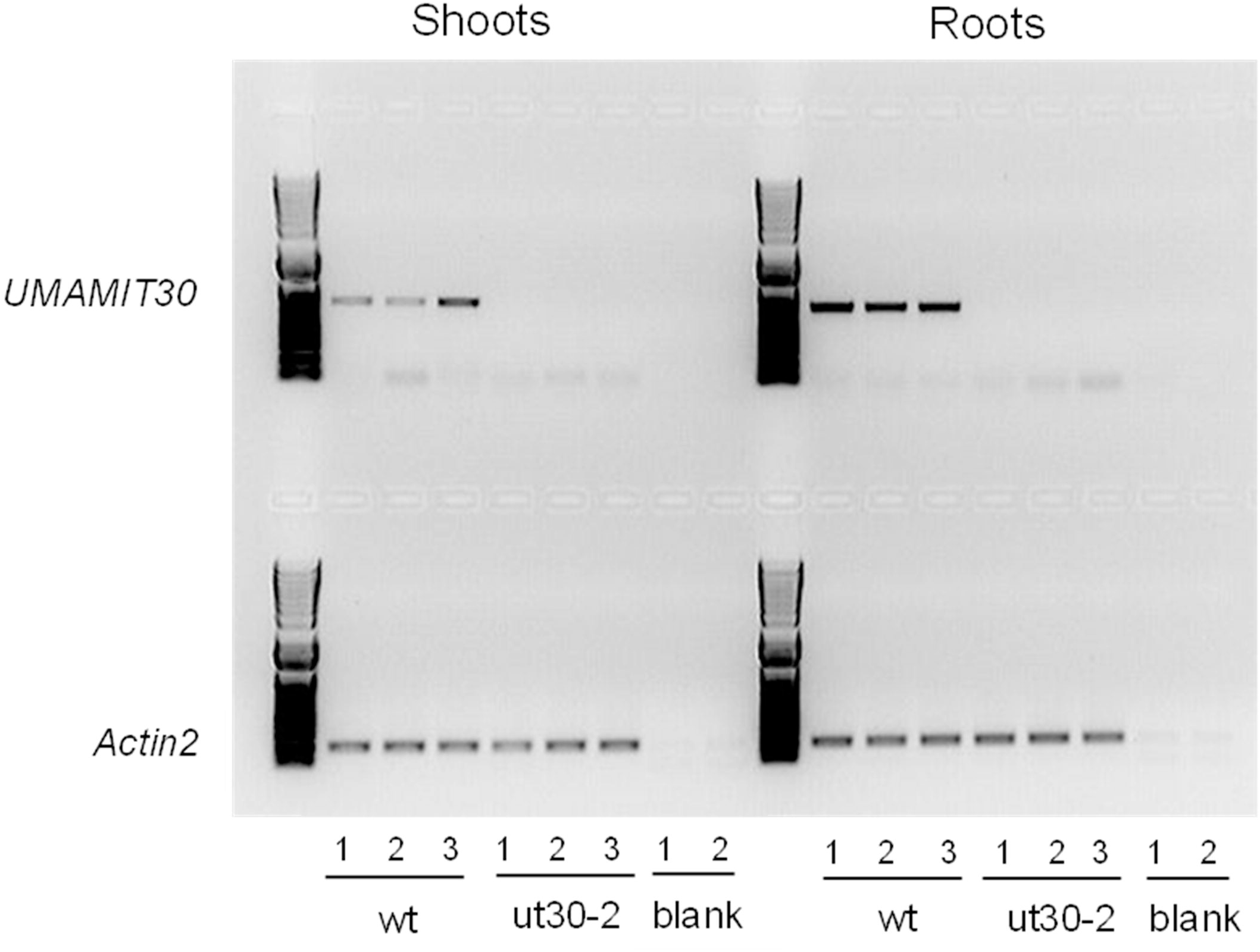
Analysis of *UMAMIT30* gene expression in wild-type and *umamit30-2* plants by RT-PCR. Blank = no template control. Source gel (uncropped full-length gel) for Figure 2B in the main text. RT-PCR primers are located in Table S1.

**Figure S3:**
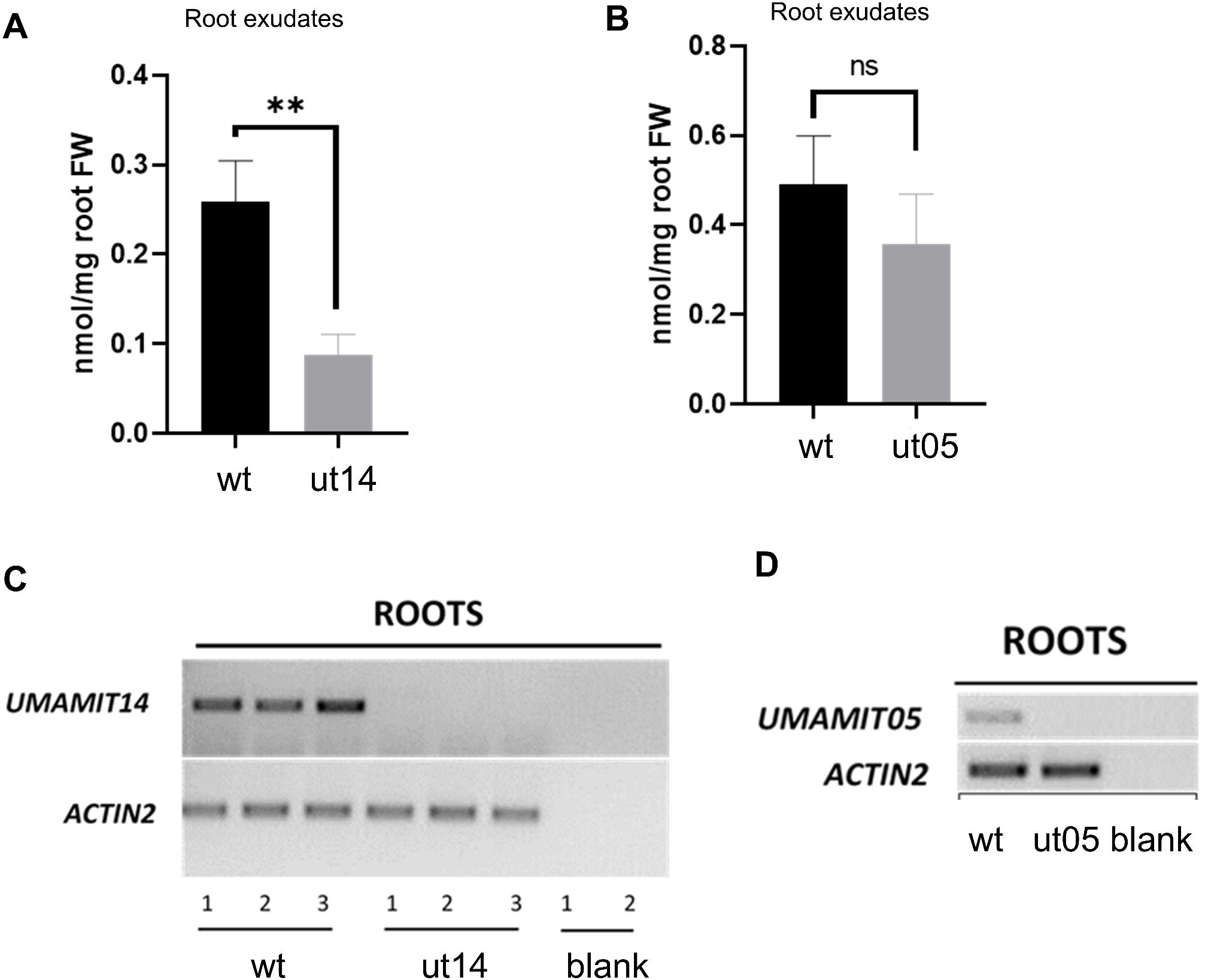
Concentrations of total free amino acids in root exudates of *umamit14* and *umamit05* mutants, compared to wild-type root exudates, and genetic analysis of mutants. Concentration of total amino acids in root exudates of wildtype vs *umamit14* plants **(A)** and wildtype vs *umamit05* plants **(B)**. AVR ± SE (*n*=6). Statistically significant difference (*p* ≤ 0.01) between wt and ut14 determined by a two- sided Student’s *t*-test is indicated by (**). ns: Non-statistically significant difference between wt and ut05. Two of two experiments yielded similar results. (C, D) *UMAMIT14* and *UMAMIT05* gene expression in wild- type, *umamit14* and *umamit05* roots of 14-day-old seedlings. *ACTIN2* expression was used as a control for cDNA input. Blank = no template control. Primers used for gene expression analyses (RT-PCR) are located in Table S1.

**Figure S4:**
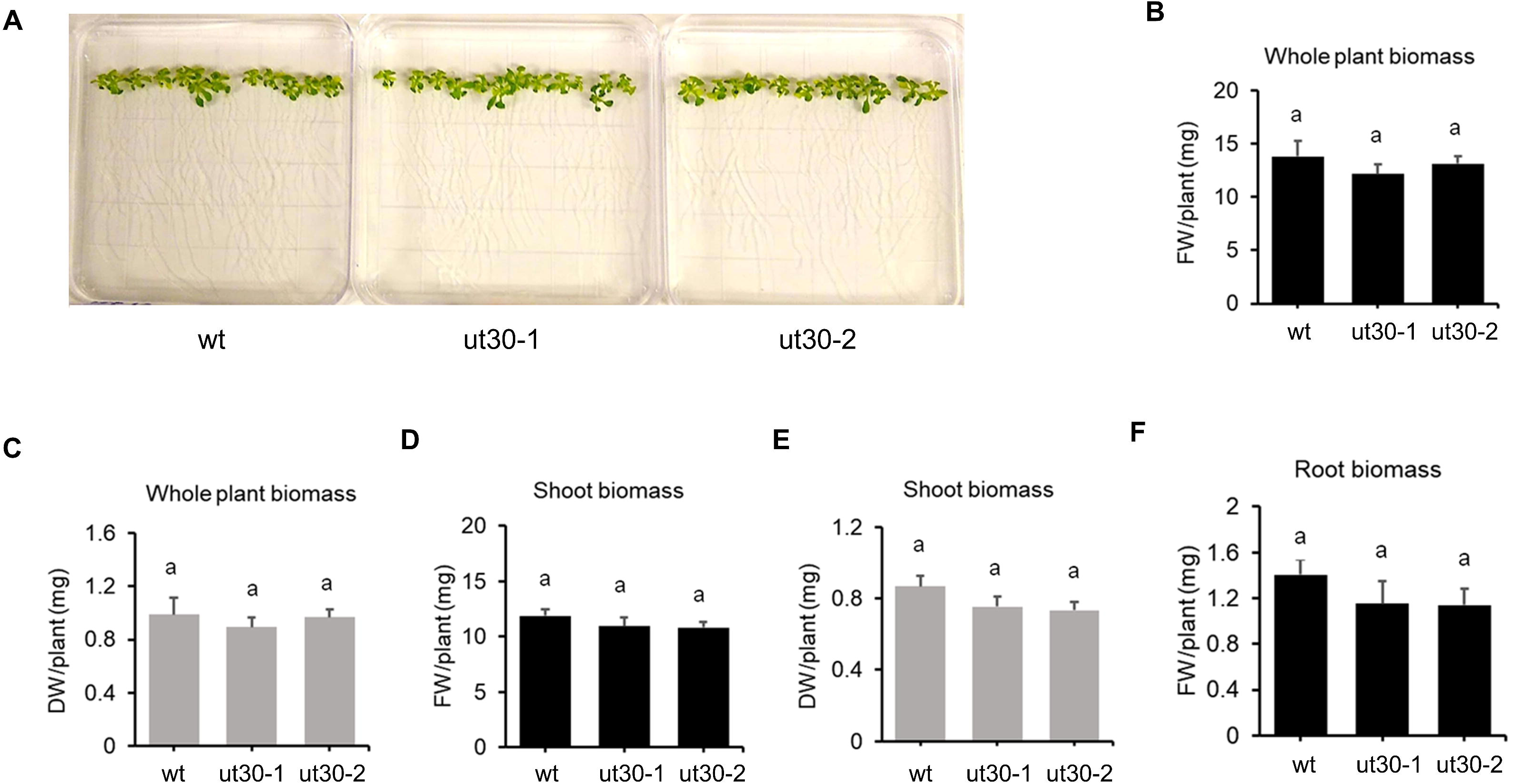
Growth phenotype analyses of wild-type and *umamit30* seedlings. **(A)** Two-week-old seedlings of wildtype, *umamit30-1*, and *umamit30-2*. Fresh (FW) and dry weights (DW) of whole plants **(B, C)** and shoots **(D, E)** of wildtype, *umamit30-1*, and *umamit30-2.* **(F)** Fresh weight of roots of wildtype, *umamit30-1*, and *umamit30-2* plants. No dry weight data is shown for roots, as they were too small to weigh accurately for individual plants. AVR ± SE (*n*=15). Bars carrying same letters indicate no statistically significant difference between the means (*p* ≤ 0.05; One-way ANOVA. Kruskal-Wallis test for unequal variances). Experiment was performed twice with similar results.

**Figure S5:**
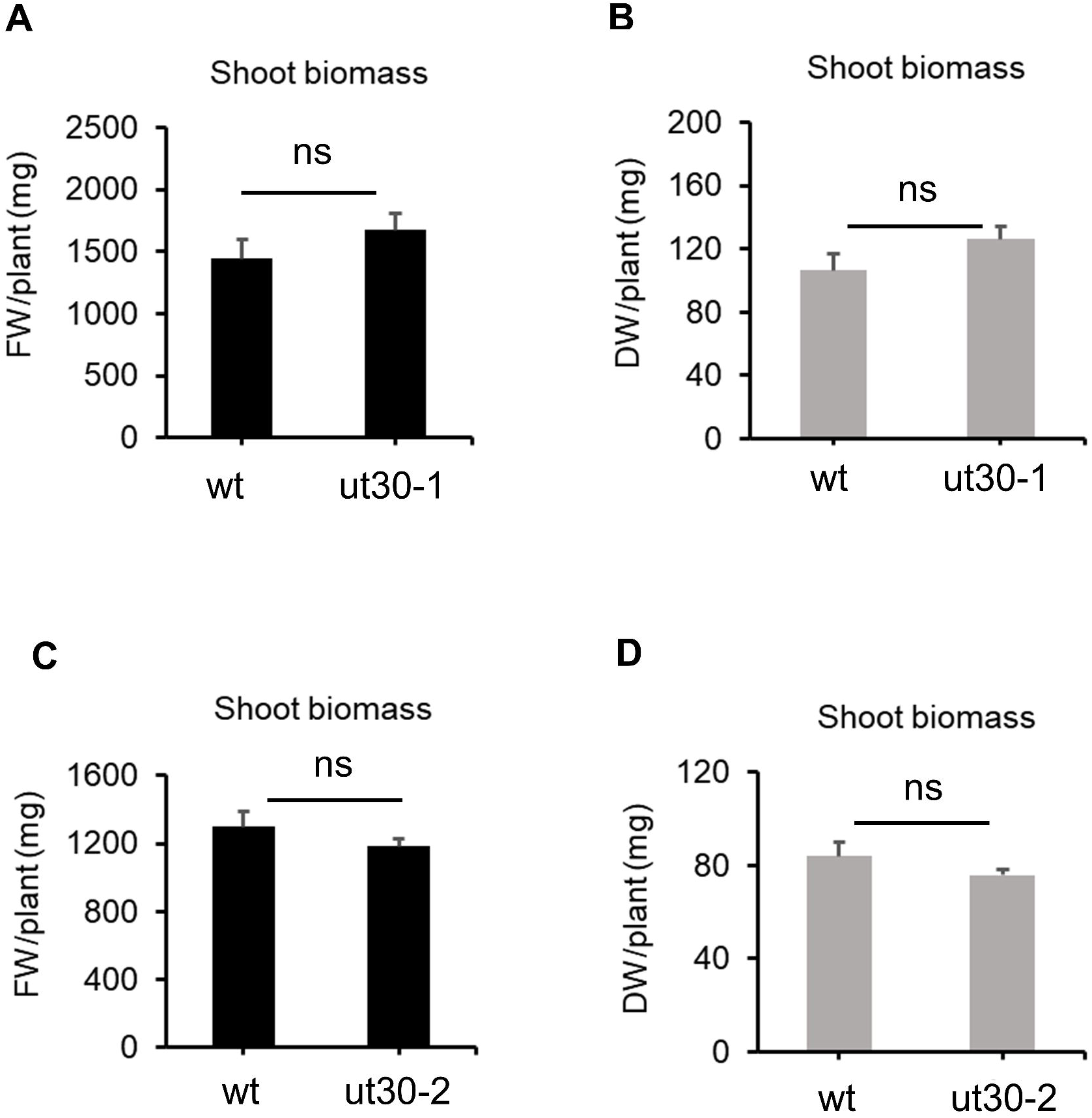
**Growth phenotype analyses of wild-type and *umamit30* mature plants.** Fresh and dry weights of shoots of 6.5-week-old wild-type and *umamit30-1* plants **(A, B)** and shoots of 6-week-old wild- type and *umamit30-2* plants **(C, D)**. AVR ± SE (*n*=10). ‘ns’ indicates no statistically significant difference between the means (*p* ≤ 0.05; two-sided Students *t*-test). Experiment was performed twice with similar results.

**Figure S6:**
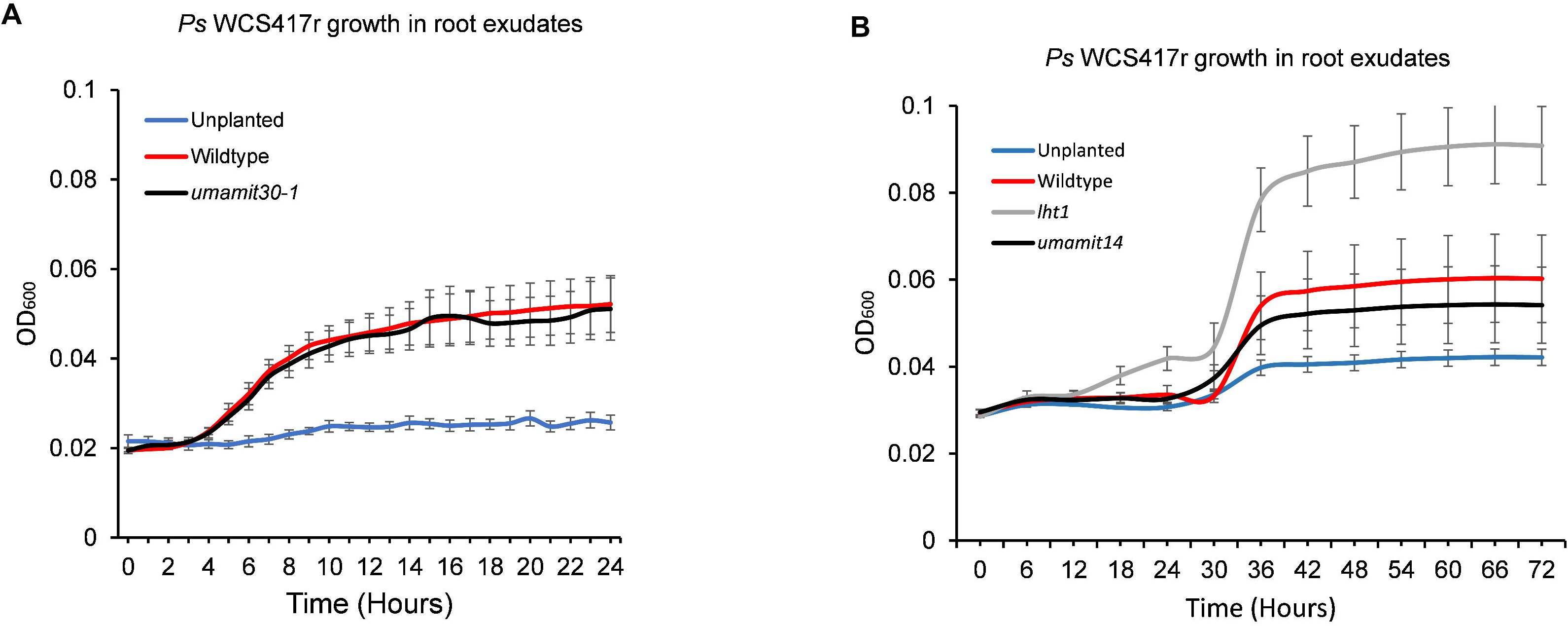
***Ps* WCS417r growth in root exudates, and in unplanted control**. **(A)** *Ps* WCS417r growth in wildtype and *umamit30* root exudates, as well as in unplanted control. **(B)** *Ps* WCS417r growth in wildtype, *lht1*, and *umamit14* root exudates, as well as in unplanted control. For (A), data are average ± SE (*n*=6 biological replicates). For (B), data are average ± SE (*n*=6 biological replicates x 2 = 12 replicates). Statistical analyses were performed using ‘Compare Groups of Growth Curves’ method as previously described [38], with the following results: (A) no significant difference between bacterial growth in wildtype and *umamit30-1* exudates. (B) wildtype vs *umamit14*: *p*=0.8719; wildtype vs *lht1*: *p*=0.0400. Experiment in (A) was performed three times with similar results.

**Table S1:**
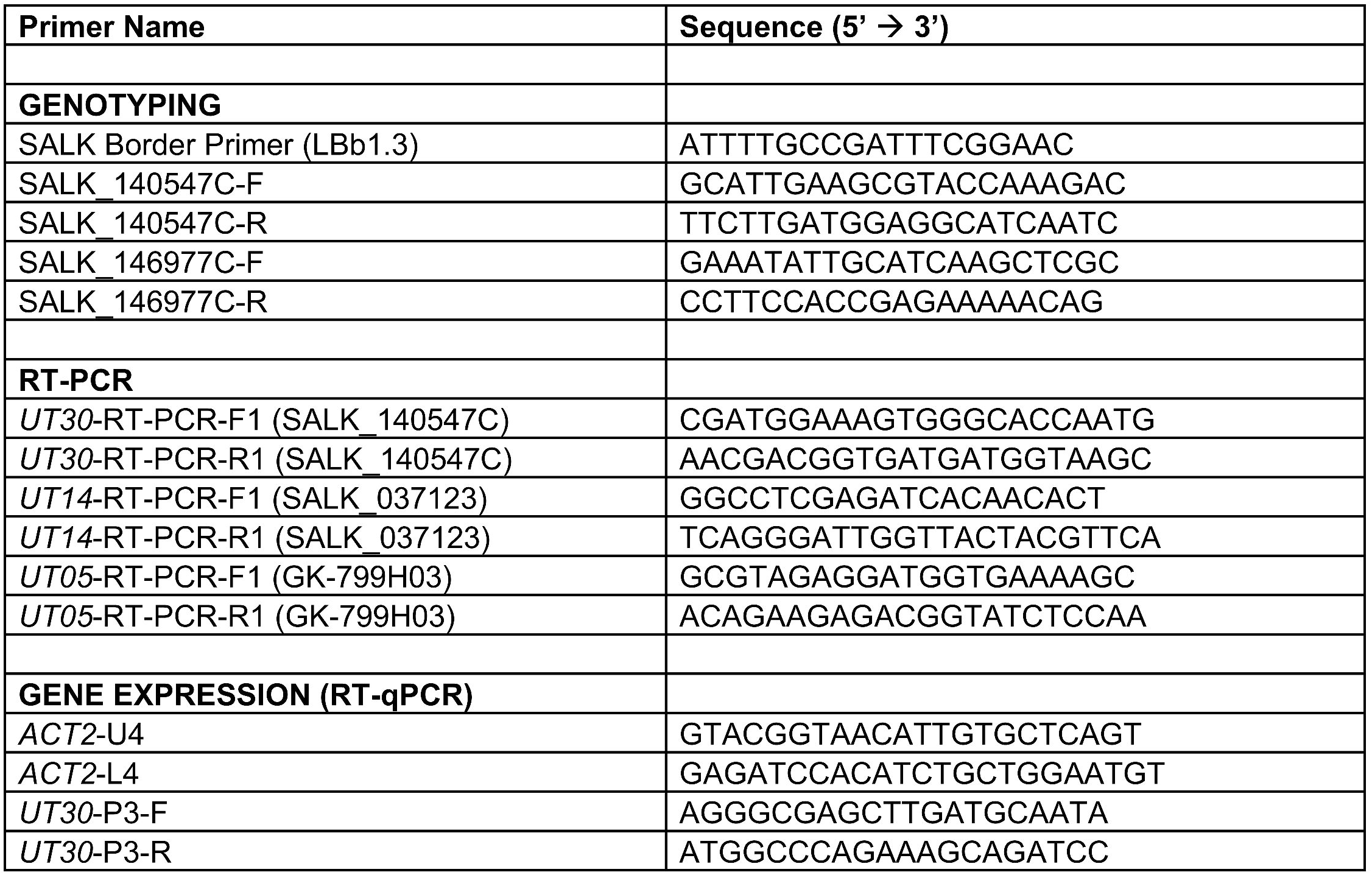
Primers used in this study.

## References

1. Sasse J, Martinoia E, Northen T. Feed your friends: do plant exudates shape the root microbiome? Trends Plant Sci. 2018;23(1):25–41. https://www.sciencedirect.com/science/article/pii/S1360138517301991

2. de la Fuente Cantó C, Simonin M, King E, Moulin L, Bennett MJ, Castrillo G, et al. An extended root phenotype: the rhizosphere, its formation and impacts on plant fitness. Plant J. 2020;103:951–64.

3. Lynch JM, Whipps JM. Substrate flow in the rhizosphere. Plant Soil. 1990;129(1):1–10.

4. Cesco S, Neumann G, Tomasi N, Pinton R, Weisskopf L. Release of plant-borne flavonoids into the rhizosphere and their role in plant nutrition. Plant Soil. 2010;329(1):1–25.

5. Woong JP, Hochholdinger F, Gierl A. Release of the benzoxazinoids defense molecules during lateral- and crown root emergence in *Zea mays*. J Plant Physiol. 2004;161(8):981–5.

6. Hartmann A, Schmid M, van Tuinen D, Berg G. Plant-driven selection of microbes. Plant Soil. 2009;321(1–2):235–57.

7. Haichar F el Z, Santaella C, Heulin T, Achouak W. Root exudates mediated interactions belowground. Soil Biol Biochem. 2014;77:69–80. 10.1016/j.soilbio.2014.06.017

8. Allard-Massicotte R, Tessier L, Lécuyer F, Lakshmanan V, Lucier JF, Garneau D, et al. *Bacillus subtilis* early colonization of *Arabidopsis thaliana* roots involves multiple chemotaxis receptors. MBio. 2016;7(6):e01664.

9. Lugtenberg BJJ, Dekkers L, Bloemberg G V. Molecular determinants of rhizosphere colonization by *Pseudomonas*. Annu Rev Phytopathol. 2001;39:461– 90.

10. Jonkers W, Rodrigues CDA, Rep M. Impaired colonization and infection of tomato roots by the Δfrpl mutant of *Fusarium oxysporum* correlates with reduced CWDE gene expression. Mol Plant-Microbe Interact. 2009;22(5):507–18.

11. Mäder U, Homuth G, Scharf C, Büttner K, Bode R, Hecker M. Transcriptome and proteome analysis of *Bacillus subtilis* gene expression modulated by amino acid availability. J Bacteriol. 2002;184(15):4288–95.

12. Moe LA. Amino acids in the rhizosphere: from plants to microbes. Am J Bot. 2013;100(9):1692–705.

13. Hoskisson PA, Sharples GP, Hobbs G. The importance of amino acids as carbon sources for *Micromonospora echinospora* (ATCC 15837). Lett Appl Microbiol. 2003;36(5):268–71.

14. Moreno R, Martínez-Gomariz M, Yuste L, Gil C, Rojo F. The *Pseudomonas putida* Crc global regulator controls the hierarchical assimilation of amino acids in a complete medium: evidence from proteomic and genomic analyses. Proteomics. 2009;9(11):2910–28.

15. Hirsch AM, Valdés M. *Micromonospora*: an important microbe for biomedicine and potentially for biocontrol and biofuels. Soil Biol Biochem. 2010;42(4):536–42.

16. Zinser ER, Kolter R. Mutations enhancing amino acid catabolism confer a growth advantage in stationary phase. J Bacteriol. 1999;181(18):5800–7.

17. Nelson KE, Weinel C, Paulsen IT, Dodson RJ, Hilbert H, Martins Dos Santos VAP, et al. Complete genome sequence and comparative analysis of the metabolically versatile *Pseudomonas putida* KT2440. Environ Microbiol. 2002;4(12):799–808.

18. Ortiz-Lopez A, Chang H-C, Bush DR. Amino acid transporters in plants. Biochim Biophys Acta. 2000;1465:275–80.

19. Chaparro JM, Badri D V., Bakker MG, Sugiyama A, Manter DK, Vivanco JM. Root exudation of phytochemicals in *Arabidopsis* follows specific patterns that are developmentally programmed and correlate with soil microbial functions. PLoS One. 2013;8(2):1–10.

20. Zhalnina K, Louie KB, Hao Z, Mansoori N, da Rocha UN, Shi S, et al. Dynamic root exudate chemistry and microbial substrate preferences drive patterns in rhizosphere microbial community assembly. Nat Microbiol. 2018;3(4):470–80. http://www.nature.com/articles/s41564-018-0129-3

21. Frommer WB, Hummel S, Unseld M, Ninnemann O. Seed and vascular expression of a high-affinity transporter for cationic amino acids in *Arabidopsis*. Proc Natl Acad Sci. 1995;92:12036–40.

22. Fischer W-N, Kwart M, Hummel S, Frommer WB. Substrate specificity and expression profile of amino acid transporters (AAPs) in *Arabidopsis*. J Biol Chem. 1995;270(27):16315–20.

23. Frommer WB, Hummel S, Riesmeier JW. Expression cloning in yeast of a cDNA encoding a broad specificity amino acid permease from *Arabidopsis thaliana*. Proc Natl Acad Sci. 1993;90:5944–8.

24. Chen L, Bush DR. LHT1, a lysine- and histidine-specific amino acid transporter in *Arabidopsis*. Plant Physiol. 1997;115(3):1127–34. http://www.ncbi.nlm.nih.gov/pubmed/9390441

25. Agorsor IDK, Kagel BT, Danna CH. The *Arabidopsis* LHT1 amino acid transporter contributes to *Pseudomonas simiae*-mediated plant growth promotion by modulating bacterial metabolism in the rhizosphere. Plants. 2023;12(2):371.

26. Okumoto S, Pilot G. Amino acid export in plants: a missing link in nitrogen cycling. Mol Plant. 2011;4(3):453–63. 10.1093/mp/ssr003

27. Ladwig F, Stahl M, Ludewig U, Hirner AA, Hammes UZ, Stadler R, et al. *Siliques Are Red1* from *Arabidopsis* acts as a bidirectional amino acid transporter that is crucial for the amino acid homeostasis of siliques. Plant Physiol. 2012;158(4):1643–55.

28. Müller B, Fastner A, Karmann J, Mansch V, Hoffmann T, Schwab W, et al. Amino acid export in developing *Arabidopsis* seeds depends on UmamiT facilitators. Curr Biol. 2015;25(23):3126–31.

29. Besnard J, Pratelli R, Zhao C, Sonawala U, Collakova E, Pilot G, et al. UMAMIT14 is an amino acid exporter involved in phloem unloading in *Arabidopsis* roots. J Exp Bot. 2016;67(22):6385–97. https://academic.oup.com/jxb/article-lookup/doi/10.1093/jxb/erw412

30. Besnard J, Zhao C, Avice JC, Vitha S, Hyodo A, Pilot G, et al. *Arabidopsis* UMAMIT24 and 25 are amino acid exporters involved in seed loading. J Exp Bot. 2018;69(21):5221–32.

31. Besnard J, Sonawala U, Maharjan B, Collakova E, Finlayson SA, Pilot G, et al. Increased expression of UMAMIT amino acid transporters results in activation of salicylic acid dependent stress response. Front Plant Sci. 2021;11:1–10.

32. Zhao C, Pratelli R, Yu S, Shelley B, Collakova E, Pilot G. Detailed characterization of the UMAMIT proteins provides insight into their evolution, amino acid transport properties, and role in the plant. J Exp Bot. 2021;72(18):6400–17.

33. The SV, Santiago JP, Pappenberger C, Hammes UZ, Tegeder M. UMAMIT44 is a key player in glutamate export from *Arabidopsis* chloroplasts. Plant Cell. 2024;36(4):1119–39. 10.1093/plcell/koad310

34. Brady S, Orlando D, JY L, Wang J, Koch J, Dinneny JR, et al. A high-resolution root spatiotemporal map reveals dominant expression patterns. Science. 2007;318:801–7.

35. Alonso JM, Stepanova AN, Leisse TJ, Kim CJ, Chen H, Shinn P, et al. Genome- wide insertional mutagenesis of *Arabidopsis thaliana*. Science. 2003;301(5633):653–7. http://science.sciencemag.org/

36. Haney CH, Samuel BS, Bush J, Ausubel FM. Associations with rhizosphere bacteria can confer an adaptive advantage to plants. Nat Plants. 2015;1(Article number: 15051). www.nature.com/natureplants

37. Lamers J, Schippers B, Geels F. Soil-borne diseases of wheat in the Netherlands and results of seed bacterization with *Pseudomonads* against *Gaeumannomyces graminis* var. *tritici*, associated with disease resistance. In: Jorna ML, Slootmaker LAJ (eds) Cereal breeding related to … 1988.

38. Elso CM, Roberts LJ, Smyth GK, Thomson RJ, Baldwin TM, Foote SJ, et al. Leishmaniasis host response loci (lmr1-3) modify disease severity through a Th1/Th2-independent pathway. Genes Immun. 2004 Mar 11;5(2):93–100.

39. Schmid M, Davison TS, Henz SR, Pape UJ, Demar M, Vingron M, et al. A gene expression map of *Arabidopsis thaliana* development. Nat Genet. 2005;37(5):501–6. http://www.nature.com/naturegenetics

40. Winter D, Vinegar B, Nahal H, Ammar R, Wilson G V., Provart NJ. An “Electronic Fluorescent Pictograph” browser for exploring and analyzing large-scale biological data sets. PLoS One. 2007;2(8):e718. https://dx.plos.org/10.1371/journal.pone.0000718

41. Goda H, Sasaki E, Akiyama K, Maruyama-Nakashita A, Nakabayashi K, Li W, et al. The AtGenExpress hormone and chemical treatment data set: experimental design, data evaluation, model data analysis and data access. Plant J. 2008;55(3):526–42. http://doi.wiley.com/10.1111/j.1365-313X.2008.03510.x

42. Kim JY, Symeonidi E, Pang TY, Denyer T, Weidauer D, Bezrutczyk M, et al. Distinct identities of leaf phloem cells revealed by single cell transcriptomics. Plant Cell. 2021;33(3):511–30.

43. Denyer T, Ma X, Klesen S, Scacchi E, Nieselt K, Timmermans MCP. Spatiotemporal developmental trajectories in the *Arabidopsis* root revealed using high-throughput single-cell RNA sequencing. Dev Cell. 2019;48(6):840–852.e5. 10.1016/j.devcel.2019.02.022

44. Ma X, Denyer T, Timmermans MCP. PscB: A browser to explore plant single cell RNA-sequencing data sets. Plant Physiol. 2020;183(2):464–7.

45. Hirner A, Ladwig F, Stransky H, Okumoto S, Keinath M, Harms A, et al. *Arabidopsis* LHT1 is a high-affinity transporter for cellular amino acid uptake in both root epidermis and leaf mesophyll. Plant Cell. 2006;18(8):1931–46. http://www.ncbi.nlm.nih.gov/pubmed/16816136

46. Hunt E, Gattolin S, Newbury HJ, Bale JS, Tseng HM, Barrett DA, et al. A mutation in amino acid permease AAP6 reduces the amino acid content of the *Arabidopsis* sieve elements but leaves aphid herbivores unaffected. J Exp Bot. 2010;61(1):55– 64.

47. Hammes UZ, Nielsen E, Honaas LA, Taylor CG, Schachtman DP. AtCAT6, a sink-tissue-localized transporter for essential amino acids in *Arabidopsis*. Plant J. 2006;48(3):414–26.

48. Elashry A, Okumoto S, Siddique S, Koch W, Kreil DP, Bohlmann H. The AAP gene family for amino acid permeases contributes to development of the cyst nematode *Heterodera schachtii* in roots of *Arabidopsis*. Plant Physiol Biochem. 2013;70:379–86. 10.1016/j.plaphy.2013.05.016

49. Pariyar SR, Nakarmi J, Anwer MA, Siddique S, Ilyas M, Elashry A, et al. Amino acid permease 6 modulates host response to cyst nematodes in wheat and Arabidopsis. Nematology. 2018;20(8):737–50.

50. Marella HH, Nielsen E, Schachtman DP, Taylor CG. The amino acid permeases AAP3 and AAP6 are involved in root-knot nematode parasitism of *Arabidopsis*. Mol Plant-Microbe Interact. 2013;26(1):44–54.

51. Dutta TK, Rupinikrishna K, Akhil VS, Vashisth N, Phani V, Pankaj, et al. CRISPR/Cas9-induced knockout of an amino acid permease gene (AAP6) reduced *Arabidopsis thaliana* susceptibility to *Meloidogyne incognita*. BMC Plant Biol. 2024;24(1):1–14.

52. Lalonde S, Tegeder M, Throne-Holst M, Frommer WB, Patrick JW. Phloem loading and unloading of sugars and amino acids. Plant, Cell Environ. 2003;26(1):37–56.

53. Tegeder M, Masclaux-Daubresse C. Source and sink mechanisms of nitrogen transport and use. New Phytol. 2018;217(1):35–53.

